# KAS-CUT&Tag for direct mapping of transcription bubbles

**DOI:** 10.64898/2026.05.15.725569

**Authors:** Weifang Wu, Jacob E. Greene, Kami Ahmad, Steven Henikoff

## Abstract

Transcription by RNA Polymerase II (Pol II) generates dynamic transcription bubbles as it moves forward, but existing methods map transcriptional activity indirectly. Here, we introduce KAS-CUT&Tag, a method that combines N_3_-kethoxal labeling of exposed single-stranded DNA with CUT&Tag to directly map transcription bubbles. Bubble density correlates with nascent transcription levels and varies across Pol II-bound genes at promoters, gene bodies, and termination regions. Transcription bubbles are enriched at genes marked by active histone modifications H3K27ac, H3K4me3, and H3K36me3 and by chromatin-bound splicing factor U2AF2, but enrichment is highest at replication-coupled histone genes in K562 and GM12878 cells. KAS-CUT&Tag also detects co-occupancy of the histone gene transcription factor NPAT with Pol II at the Histone Locus Body, suggesting that in these cells NPAT is juxtaposed to transcription bubbles via interaction with Pol II. Together, KAS-CUT&Tag enables direct mapping of Pol II and regulatory interactions at transcription bubbles, providing a powerful tool for precise analysis of transcriptional activity.

## Introduction

RNA polymerase II (Pol II) transcribes all protein-coding genes and many non-coding RNAs in the eukaryotic nucleus. Transcription begins with recruitment of unphosphorylated Pol II to promoters, forming a closed Pre-Initiation Complex (PIC) with general transcription factors. Transcription initiates with DNA unwinding at the promoter to form an open PIC with a single-stranded transcription bubble, facilitated by TFIIH and other general transcription factors^1–4^. Following transcriptional initiation, Pol II is phosphorylated on serine-5 (Pol II-Ser5p) of the largest subunit RPB1 C-Terminal Domain (CTD) heptapeptide repeats^5–7^ and then moves forward, but often pauses ∼20–60 nucleotides downstream of the transcription start site (TSS)^8,9^. Release of this paused complex into active elongation is regulated by CDK9-mediated phosphorylation of the Pol II CTD at serine-2 (Pol II-Ser2p)^10,11^. After Pol II proceeds through the body of genes, transcription is terminated and Pol II disengages from its DNA template, whereupon the transcription bubble reanneals^12,13^.

Pol II translocates the single-stranded transcription bubble as it moves forward. Within this transcription bubble, the nascent RNA pairs with the template DNA as an RNA–DNA hybrid, positioning the RNA 3′ end in the Pol II catalytic site, while the non-template single-stranded DNA (ssDNA) is displaced^14,15^. As Pol II translocates, it exposes the next template base for pairing with an incoming nucleoside triphosphate, extending the RNA by one nucleotide^16^. Thus, the transcription bubble moves dynamically as Pol II progresses.

To map these ssDNA structures, we developed KAS-CUT&Tag, a method that combines N_3_-kethoxal labeling of exposed guanines in single-stranded bubbles with CUT&Tag. Applied to Pol II, KAS-CUT&Tag maps transcription bubbles at Pol II-bound sequences across the genome and shows that bubble density varies within and between loci. Perturbation experiments confirm that these bubbles arise from actively engaged Pol II. Pol II is highly enriched at replication-coupled histone genes and precisely co-localizes with the transcription factor NPAT in K562 cells. We conclude that KAS-CUT&Tag enables high-resolution mapping of transcription bubbles and anticipate broad applications in dissecting transcriptional regulatory mechanisms.

## Results

### KAS-CUT&Tag detects single-stranded DNA bound by Pol II

To efficiently map single-stranded transcription bubbles, we capitalized on N_3_-kethoxal-assisted single-stranded DNA sequencing (KAS-seq), where reaction of unpaired guanine nucleosides with N_3_-kethoxal is followed by biotin labeling and immunoprecipitation^17^. Cells are immobilized on Concanavalin A (ConA) beads and permeabilized, followed by a 10-minute treatment with N_3_-kethoxal. We then proceeded with antibody-tethered Tn5 tagmentation, fragment release, and DNA purification. This DNA is then biotinylated via click chemistry, and biotinylated DNA is immunoprecipitated with streptavidin beads and subjected to PCR for library construction. We call this general method kethoxal-assisted single-stranded DNA assay for CUT&Tag (KAS-CUT&Tag) (Fig. 1a). Parallel reactions omitting the immunoprecipitation step serve as KAS-CUT&Tag total library profiles.

**Fig. 1:**
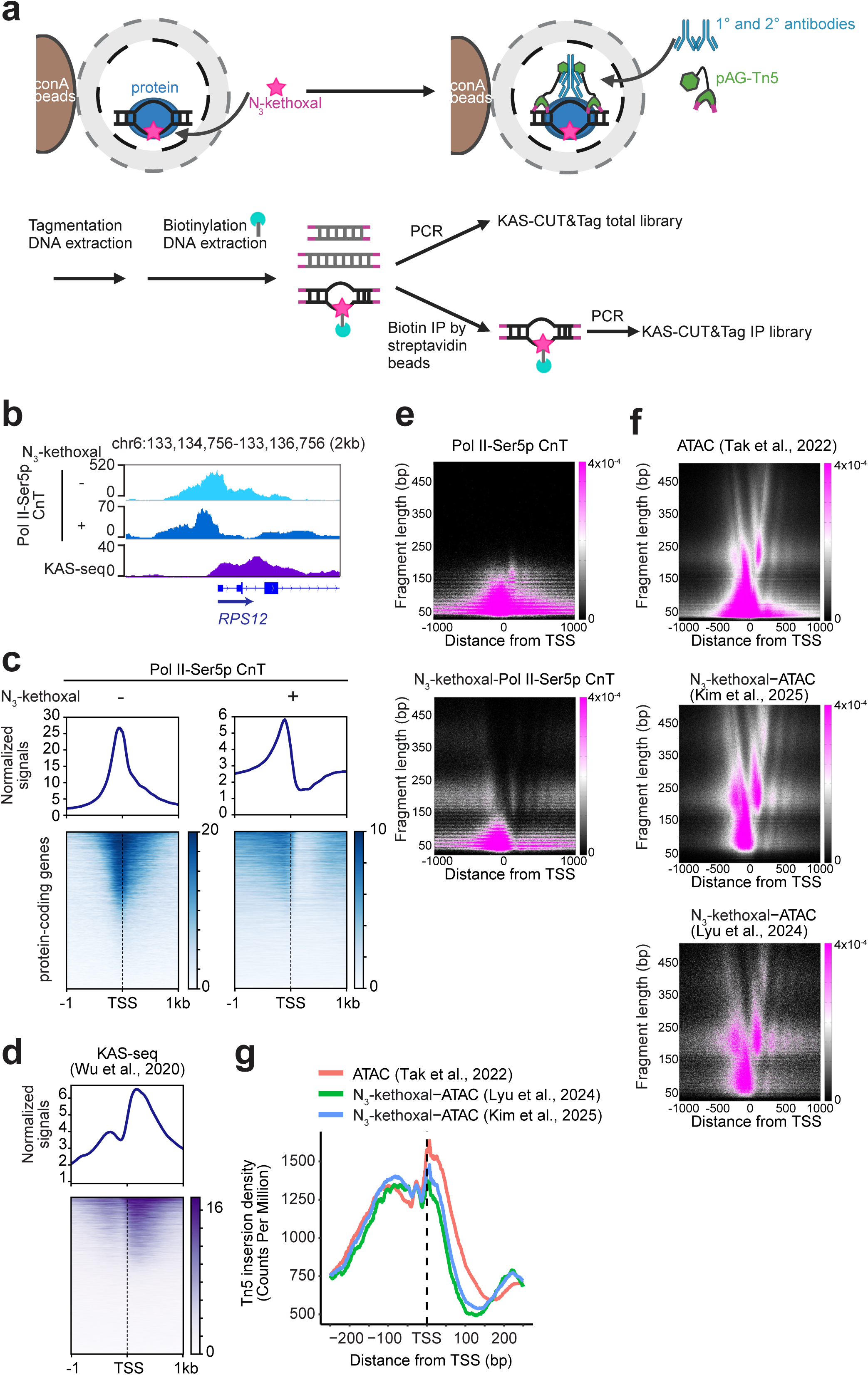
KAS-CUT&Tag detects single-stranded DNA bound by Pol II. **a,** Schematic of the KAS-CUT&Tag workflow (Created in BioRender. Wu, W. (2026) https://BioRender.com/2ffxgwv) N_3_-kethoxal (magenta star) labels ssDNA, while a primary antibody (blue Y-shape) binds the target protein (blue oval). A secondary antibody (blue Y-shape) enhances recruitment of the pAG-Tn5 transposome (green), which is activated by Mg²⁺ to insert adapters (magenta lines) at protein-bound sites. After DNA purification, adapter-ligated fragments are biotinylated via click chemistry. Biotinylated DNA is either amplified by PCR to generate the KAS-CUT&Tag (KAS-CnT) total library, or immunoprecipitated with streptavidin beads before PCR to construct the KAS-CnT IP library. **b,** Coverage-normalized Pol II-Ser5p CUT&Tag (CnT) signals (±N_3_-kethoxal), shown alongside KAS-seq data from Wu et al in K562 cells^17^. KAS-CnT total library corresponds to N_3_-kethoxal-treated CnT. **c, d,** Heatmaps (bottom) and average plots (top) centered on the TSSs of 12,397 protein-coding genes, showing Pol II-Ser5p CnT (±N_3_-kethoxal) (**c**) and KAS-seq data in K562 (**d**). Each row represents one gene. **e, f,** V-plots showing fragment size versus distance from TSSs for Pol II-Ser5p CnT (±N_3_-kethoxal) (**e**) and ATAC-seq (±N_3_-kethoxal) (**f**). ATAC-seq (–N_3_-kethoxal) is from Tak et al^20^, while ATAC-seq (+N_3_-kethoxal) is from Kim et al^21^ and Lyu et al^22^. **g,** Density plot of Tn5 fragment ends within a 400 bp window centered on TSSs.

We first applied KAS-CUT&Tag using antibodies against Pol II and Ser2-phosphorylated Pol II in human K562 cells. We optimized the immunoprecipitation step, yielding highly specific and reproducible KAS-CUT&Tag profiles (Extended Data Fig.1a-d). We used streptavidin T1 beads for immunoprecipitation in all subsequent KAS-CUT&Tag experiments. KAS-CUT&Tag IP profiles correlated with total controls over gene bodies but not with GC content (Extended Data Fig.1e,f), confirming that the labeling method was not biased. Because light crosslinking with 0.1% formaldehyde and low-salt CUTAC tagmentation enabled single-cell Pol II profiling in our previously published CUT&Tag2for1 and sci-CUT&Tag2in1 methods^18,19^, we reasoned that modifying KAS-CUT&Tag to include these conditions could improve library preparation. Light crosslinking after N_3_-kethoxal treatment followed by low-salt CUTAC tagmentation increased KAS-CUT&Tag library yield, while maintaining high reproducibility and profile qualities (Extended Data Fig.2a–d).

We next examined whether N_3_-kethoxal treatment affects CUT&Tag profiles by analyzing KAS-CUT&Tag total libraries, which represent N_3_-kethoxal-treated CUT&Tag. Compared to untreated CUT&Tag, N_3_-kethoxal-treated CUT&Tag for Pol II-Ser5p showed signal depletion from the TSS to ∼+500 bp (Fig. 1b,c). Within this gap, KAS-seq ssDNA signals were enriched (Fig. 1b,d), thus N_3_-kethoxal labeling of unpaired guanines in ssDNA appears to prevent reannealing, thereby blocking Tn5 tagmentation in this region and producing the observed loss of signal. V-plots supported this interpretation: Untreated Pol II-Ser5p CUT&Tag showed dense subnucleosomal fragments around the TSS, whereas N_3_-kethoxal-treated CUT&Tag displayed a pronounced loss of subnucleosomal fragments and only a modest increase in mononucleosomal fragments downstream, consistent with the observed gap (Fig. 1e). Similarly, N3-kethoxal-treated Pol II CUT&Tag showed a reduction in subnucleosomal fragments downstream of TSSs relative to untreated Pol II CUT&Tag (Figure S3a). To test whether blocking of Tn5 tagmentation by N_3_-kethoxal is a general limitation, we compared untreated ATAC-seq^20^ and N_3_-kethoxal-treated ATAC-seq (KAS-ATAC total libraries) from published data^21,22^. In contrast to CUT&Tag, where Tn5 integration is tethered to antibody-bound chromatin, ATAC-seq utilizes freely diffusing Tn5 that targets accessible double-stranded DNA (dsDNA). V-plots of N_3_-kethoxal-treated ATAC-seq data from two different laboratories^21,22^ revealed a loss of subnucleosomal fragments around the TSS (Fig. 1f). In addition, insertion-site analysis showed marked depletion downstream of the TSS (Fig. 1g), consistent with Tn5 exclusion from ssDNA in this region. These analyses show that N_3_-kethoxal labeling creates a technical “blind spot” in both KAS-CUT&Tag and KAS-ATAC by blocking Tn5 integration into labeled ssDNA between the TSS and the +1 nucleosome (Extended Data Fig.3b). This effect, which had not previously been recognized in KAS-ATAC, leads to a loss of short fragments while preserving longer fragments derived from adjacent dsDNA. Despite this technical blind spot, efficient N_3_-kethoxal labeling of ssDNA provides the basis for KAS-CUT&Tag profiling of Pol II-associated ssDNA generated during transcription.

### KAS-CUT&Tag maps transcription bubbles

Pol II-targeted KAS-CUT&Tag total – which omits the immunoprecipitation step – profiles chromatin-bound Pol II, while Pol II-targeted KAS-CUT&Tag IP captures transcription bubbles. We used the ratio of KAS-CUT&Tag IP to KAS-CUT&Tag total signal as a measure of transcription bubble density, with higher values indicating a greater fraction of Pol II localized in transcriptional bubbles. Surprisingly, we found that bubble density varied substantially around gene TSSs bound by Pol II. For example, Pol II occupancy is similar around the TSSs of the *MYL11* and *TXNIP* genes, but both bubble density and nascent transcription measured by public PRO-seq data^23^ were markedly higher at *TXNIP* (Fig. 2a). This implies that at genes such as *MYL11*, only a fraction of chromatin-bound Pol II is in an open complex.

**Fig. 2:**
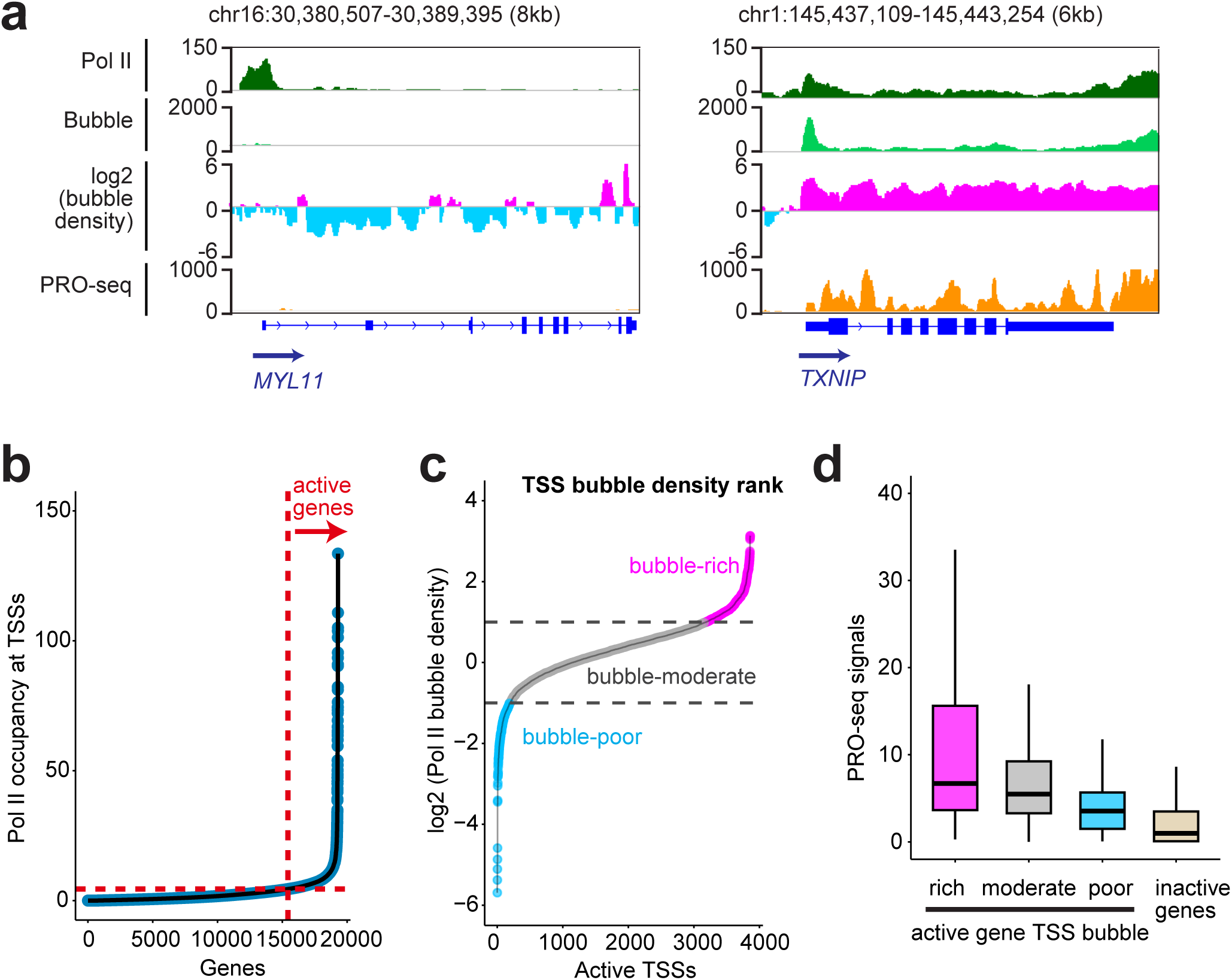
Transcription bubble density varies between gene promoters. **a,** Coverage-normalized Pol II KAS-CnT total signal, Pol II KAS-CnT IP signal corresponding to transcription bubbles, log₂(bubble density) calculated as log₂(IP/total), and PRO-seq signal in K562 cells. Nascent transcription levels were measured by public PRO-seq data^23^. **b,** Ranking of 19,294 protein-coding gene TSSs by Pol II occupancy quantified by Pol II KAS-CnT total signal in K562 cells. We classified genes whose TSSs ranked in the top 20% by Pol II occupancy as active and the remaining genes as inactive. **c,** Ranking of 3,859 active gene TSSs by log₂(bubble density) in K562. **d,** PRO-seq signals at inactive genes and active genes with bubble-rich, bubble-moderate, or bubble-poor TSSs in K562 cells. PRO-seq signals in rich (*n* = 674), moderate (*n* = 2,991), and poor (*n* = 194) active gene TSS bubble groups and inactive gene TSSs (*n* = 14,941); medians were 6.70, 5.49, 3.55, and 0.982, respectively. All pairwise comparisons were significant (two-sided Wilcoxon rank-sum tests, Benjamini–Hochberg-adjusted *p* ≤ 3.10 × 10⁻^11^). Boxes span the 25th–75th percentiles, centre lines show medians, and whiskers extend to the minimum and maximum values within 1.5× the interquartile range; outliers are not shown.

To systematically identify genes with high or low bubble density around their TSSs, we analyzed 3,859 active genes with high Pol II occupancy within a 2-kb window centered on their TSSs (Fig. 2b). Among these genes, 17% (674 TSSs) are bubble-rich (log_2_ bubble density > 1), while 5% (194 TSSs) are bubble-poor (log_2_ bubble density < –1) in K562 cells (Fig. 2c). For instance, bubble-rich *SMIM30*, bubble-moderate *GGPS1* and bubble-poor *LY6G6F* gene TSSs all showed similar Pol II occupancy (Extended Data Fig.4a). This indicates that the proportion of promoter-bound Pol II localized in an open complex varies between active genes. To directly assess the relationship between transcription and bubble density, we used PRO-seq signals to measure productive transcription. Active genes with bubble-rich TSSs showed higher PRO-seq signals than bubble-poor TSSs (Fig. 2d), suggesting that bubble density is associated with nascent transcription. Because R-loops also contain displaced ssDNA and can form during Pol II transcription, we compared bubble density with R-loop regions identified in publicly available MapR^24^, R-loop CUT&RUN^24^, and R-loop CUT&Tag^25^ datasets from HEK293T cells. R-loops were more frequently detected at active than inactive TSSs but did not correlate with bubble density among active TSSs (Extended Data Fig.4b). These results demonstrate that Pol II KAS-CUT&Tag specifically detects transcription bubbles rather than R-loops.

Active TSSs are commonly marked by histone H3 lysine 4 trimethylation (H3K4me3) and histone H3 lysine 27 acetylation (H3K27ac), whereas many inactive TSSs are marked by the repressive modification H3K27me3. Among active TSSs, bubble-rich TSSs showed higher H3K4me3 and H3K27ac but lower H3K27me3 than bubble-poor TSSs (Extended Data Fig.4c–e). Histone H3 lysine 9 trimethylation (H3K9me3), a constitutive heterochromatin-associated repressive histone modification, remained low across all TSSs (Extended Data Fig.4f). Together, these analyses show that transcription bubbles are enriched at promoters with active chromatin modifications.

We hypothesized that variation in promoter-proximal Pol II pausing contributes to differences in bubble density, with paused Pol II exposing ssDNA that is reactive to N_3_-kethoxal. To test this, we treated K562 cells with transcription inhibitors that target distinct stages of the transcription cycle (Fig. 3a). Triptolide (TPL) inhibits transcription initiation by targeting the XPB subunit of TFIIH, which catalyzes promoter opening^26^. Flavopiridol (FP) blocks pause release by inhibiting cyclin-dependent kinases^27^, including CDK9, which phosphorylates Pol II CTD serine-2. Actinomycin D (ActD) intercalates into DNA and stalls Pol II elongation^28^. We validated drug action by analyzing Pol II occupancy using KAS-CUT&Tag total signals. As expected, TPL treatment led to a global reduction in Pol II occupancy due to impaired loading (Fig. 3b; Extended Data Fig.4g); this was accompanied by a corresponding decrease in bubble density (Fig. 3c; Extended Data Fig.4h). In contrast, FP and ActD treatments increased Pol II occupancy at TSSs, consistent with blocked pause release by FP and Pol II stalling by ActD (Fig. 3b; Extended Data Fig.4g). Both treatments also significantly increased Pol II bubble density at TSSs (Fig. 3c; Extended Data Fig.4h). These results confirm that KAS-CUT&Tag detects transcription bubbles caused by Pol II pausing.

**Fig. 3:**
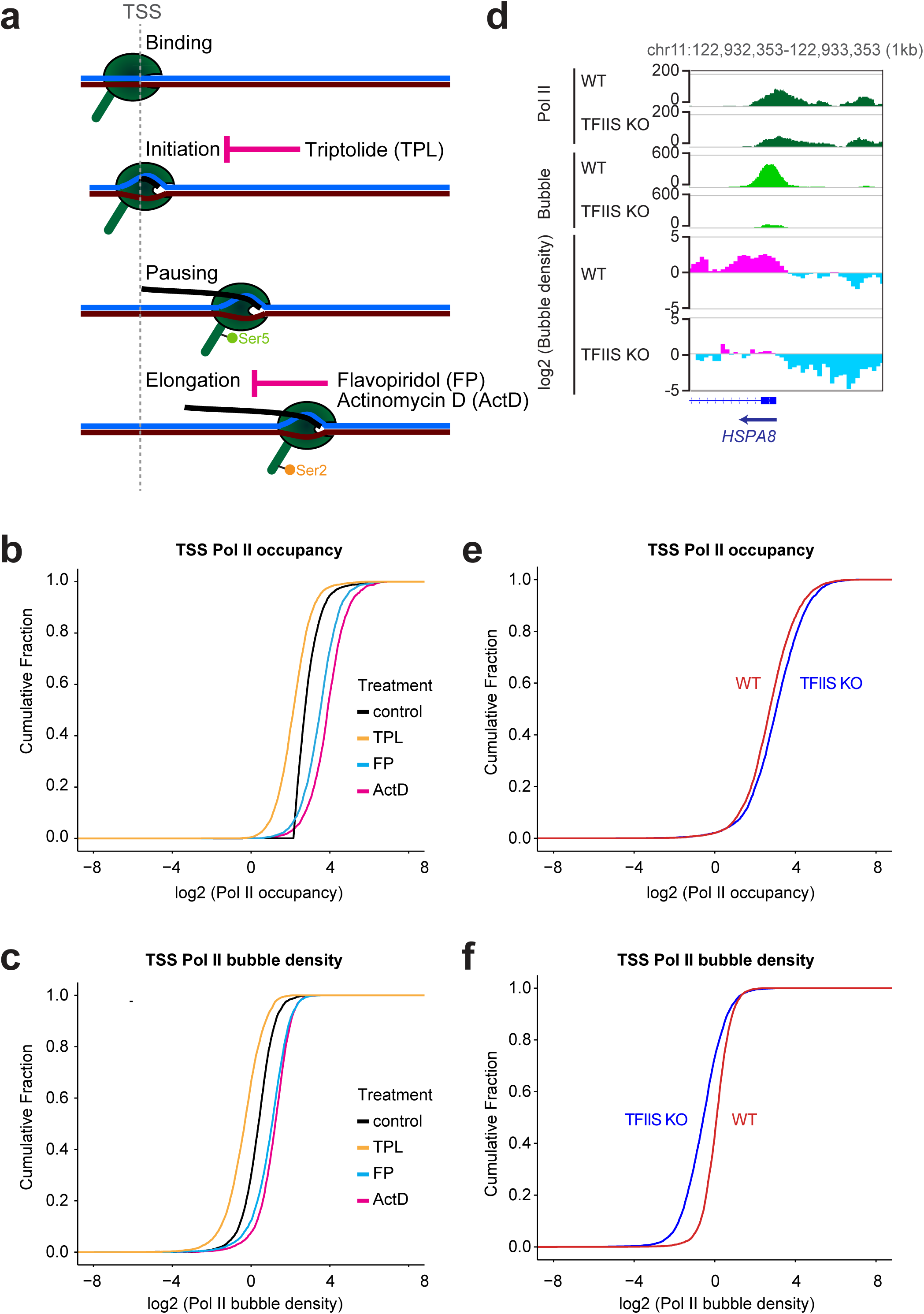
KAS-CUT&Tag maps transcription bubbles generated by Pol II. **a,** Diagram illustrating stages of the transcription cycle targeted by different transcription inhibitors. **b,** Cumulative plots showing Pol II occupancy at TSSs in K562 cells under the indicated treatments, quantified by Pol II KAS-CnT total signals. **c,** Cumulative plots showing Pol II bubble density at TSSs under the same conditions, quantified by log₂(IP/total). Two-sided Kolmogorov–Smirnov tests with Holm correction showed significant differences between each inhibitor and control. In **b**, adjusted *P* < 2.2 × 10⁻^308^ for TPL, *P* = 4.393 × 10⁻^282^ for FP and *P* < 2.2 × 10⁻^308^ for ActD; in **c**, adjusted *P* = 1.039 × 10⁻^224^ for TPL, *P* = 5.229 × 10⁻^231^ for FP and *P* < 2.2 × 10⁻^308^ for ActD. **d,** Coverage-normalized Pol II KAS-CnT total signal, Pol II KAS-CnT IP signal corresponding to transcription bubbles, and log₂(bubble density), calculated as log₂(IP/total), in WT and TFIIS KO HEK293T cells. **e,** Cumulative plots showing Pol II occupancy at TSSs in WT and TFIIS KO cells, quantified by Pol II KAS-CnT total signals. **f,** Cumulative plots showing Pol II bubble density at TSSs in WT and TFIIS KO cells, quantified by log₂(IP/total). Two-sided Kolmogorov–Smirnov tests showed significant differences between WT and TFIIS KO distributions for both Pol II occupancy (**e**, *P* = 4.99 × 10^-23^) and Pol II bubble density (**f**, *P* = 1.35 × 10^-219^).

To better understand how KAS-CUT&Tag detects transcriptionally active Pol II, we examined Pol II backtracking, a state in which the RNA 3′ end is displaced from the catalytic site^29–31^. The elongation factor TFIIS reactivates backtracked Pol II by promoting cleavage of the extruded RNA, allowing realignment of the 3′ end^29–31^. We profiled wild-type (WT) and TFIIS knockout (KO) HEK293T cells. If transcription bubbles arise from active Pol II, loss of TFIIS should trap Pol II in an inactive state and reduce bubble density without altering total occupancy. Pol II is known to backtrack at heat shock genes and requires TFIIS for reactivation^32–34^. Accordingly, the promoter of the *HSPA8* heat shock gene showed high bubble density in WT cells, but this signal was reduced in TFIIS KO cells despite only moderate change in Pol II occupancy (Fig. 3d). Genome-wide analysis confirmed that bubble density at TSSs broadly declines despite increased Pol II occupancy (Fig.3e,f). These findings demonstrate that KAS-CUT&Tag enables mapping of transcription bubbles and that TFIIS maintains these bubbles.

### Transcription bubble density varies across gene bodies and termination sites

Pol II escapes promoter pausing upon serine-2 phosphorylation of its CTD repeats and subsequently elongates across gene bodies^10,11^. We quantified Pol II-Ser2p occupancy and transcription bubbles using coverage-normalized Pol II-Ser2p KAS-CUT&Tag total and IP signals, respectively, and measured bubble density using the ratio of PoI lI-Ser2p KAS-CUT&Tag IP to KAS-CUT&Tag total signals. In contrast to the incompatibility between N_3_-kethoxal treatment and Tn5 tagmentation around TSSs, we observed minimal impact of N_3_-kethoxal on KAS-CUT&Tag total profiles of elongating Pol II-Ser2p, which remained largely unchanged across gene bodies (Extended Data Fig.3c,d).

Interestingly, as we observed at TSSs, transcription bubble densities varied substantially between active genes. For example, although the *MCL1* and *MRPL40* genes had similar Pol II-Ser2p occupancy across their gene bodies, *MCL1* had markedly higher bubble density (Fig. 4a). To systematically assess these gene body patterns while avoiding promoter-proximal signals, we quantified Pol II-Ser2p bubble density across 3,754 active genes ≥1.5 kb in length, considering the gene body as the region from +1 kb downstream of the TSS to the transcription end site (TES). Only 9% (342) of gene bodies were classified as bubble-rich, compared to 17% of active TSSs in K562 cells (Extended Data Fig.5a; Fig. 2c). Genes with bubble-rich gene bodies showed higher nascent transcription levels than genes with bubble-poor gene bodies (Extended Data Fig.5b). As expected, inhibiting CDK9-dependent pause release with flavopiridol reduced both Pol II-Ser2p occupancy and bubble density in gene bodies (Extended Data Fig.5c,d). We found that Pol II-Ser2p occupancy in TFIIS KO cells remained similar to WT cells at the *EIF4A1* gene, but bubble density was reduced (Fig. 4b). Genome-wide analysis confirmed this result: while Pol II-Ser2p occupancy was unchanged, Pol II-Ser2p bubble density was significantly reduced across gene bodies in TFIIS KO cells (Fig. 4c,d). These results suggest that backtracking resolution by TFIIS maintains Pol II-Ser2p at bubbles across gene bodies.

**Fig. 4:**
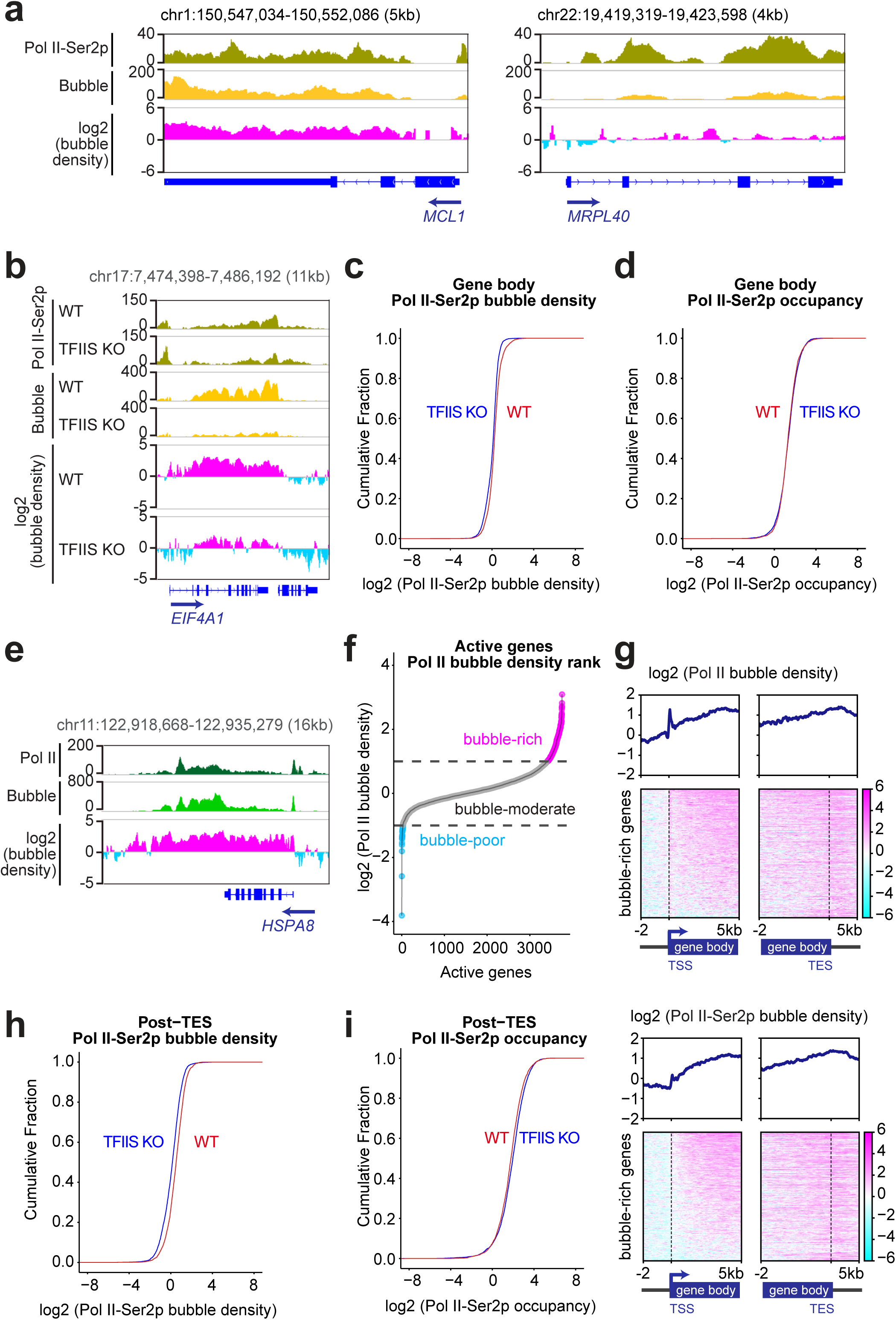
Bubble density varies across gene bodies and termination sites. **a,** Coverage-normalized Pol II-Ser2p KAS-CnT total signal, Pol II-Ser2p KAS-CnT IP signal corresponding to transcription bubbles, and log₂(bubble density), calculated as log₂(IP/total), in K562 cells. **b,** Coverage-normalized Pol II-Ser2p KAS-CnT total and IP signals, and log₂(IP/total) ratios in WT and TFIIS KO cells. **c, d,** Cumulative distribution plots showing Pol II-Ser2p bubble density (**c**) or occupancy (**d**) across gene bodies in WT and TFIIS KO cells. Bubble density was calculated as IP/total ratio, whereas occupancy was quantified using the Pol II-Ser2p KAS-CnT total signal. Two-sided Kolmogorov–Smirnov tests showed that the Pol II-Ser2p bubble density distributions differed significantly between WT and TFIIS KO cells in **c**, with *p* < 9.99 × 10^-30^, whereas no significant difference in Pol II-Ser2p occupancy was detected in **d**, with *p* = 0.062. **e,** Coverage-normalized Pol II KAS-CnT total signal, Pol II KAS-CnT IP signal, and log₂(IP/total) ratios in K562 cells. **f,** Ranking of 3,754 active genes by Pol II bubble density across entire transcription units (TSS to 2 kb downstream of TES), quantified by log₂(IP/total). Active genes were defined as genes with active TSSs, as described in Fig. 2b, and an annotated gene length greater than 1.5 kb. **g,** Heatmaps (bottom) and average plots (top) aligned to the TSSs and TESs of 343 active genes in K562, showing log₂(Pol II bubble density) or log₂(Pol II-Ser2p bubble density). Each row represents one gene. **h,i,** Cumulative plots showing Pol II-Ser2p bubble density (**h**) or occupancy (**i**) across post-TES regions in WT and TFIIS KO cells. Two-sided Kolmogorov–Smirnov tests showed significant differences between WT and TFIIS KO distributions for both bubble density (**h**, *P* = 3.5 × 10^-51^) and occupancy (**i**, *P* = 1.15 × 10^-10^).

We analyzed transcription bubble density across full transcription units, from the TSS to 2 kb downstream of the transcription end site (TES). Both Pol II and Pol II-Ser2p transcription bubble densities correlated more strongly with PRO-seq than with steady-state RNA-seq in K562 cells (Extended Data Fig.5e). We next assessed whether bubble density remains uniform across the full transcription unit. Pol II bubble density across the *HSPA8* heat shock gene was consistent from TSS to the TES in wild-type HEK293T cells (Fig. 4e). To assess this genome-wide, we used Pol II-targeted KAS-CUT&Tag total profiles to measure chromatin-bound Pol II and calculated bubble density across full transcription units. We found that 9% (343 of 3,754) of active genes were bubble-rich throughout the entire transcription unit (Fig. 4f). At these genes, Pol II and Pol II-Ser2p bubble densities were elevated at the TSS and remained relatively stable through gene bodies and post-TES regions (Fig. 4g). These findings reveal a subset of genes with uniformly high bubble density across their entire transcribed regions.

Next, we examined transcription beyond annotated TESs, where Pol II often continues elongation despite cleavage and polyadenylation. We defined the 2-kb region downstream of TESs as post-polyadenylation regions and found that 37% (1,382) were bubble-rich in K562 cells (Extended Data Fig.5f). In TFIIS KO cells, bubble density declined in these regions without a corresponding reduction in Pol II-Ser2p occupancy (Fig. 4h,i), indicating that TFIIS maintains Pol II-Ser2p at transcription bubbles downstream of TESs. These findings highlight widespread heterogeneity in bubble density at Pol II-Ser2p transcribed genes.

### Transcription bubbles are enriched at genes marked with the H3K36me3 histone modification and the U2AF2 splicing factor

Highly active gene bodies are often marked with histone H3 lysine 36 trimethylation (H3K36me3)^35,36^. To test if this modification might be associated with actively engaged Pol II, we compared the distributions of the H3K36me3 mark with bubble density generated by Pol II-Ser2p. Indeed, the H3K36me3 modification was elevated at bubble-rich gene bodies in both K562 and HEK293 cells (Fig. 5a; Extended Data Fig.6a).

**Fig. 5:**
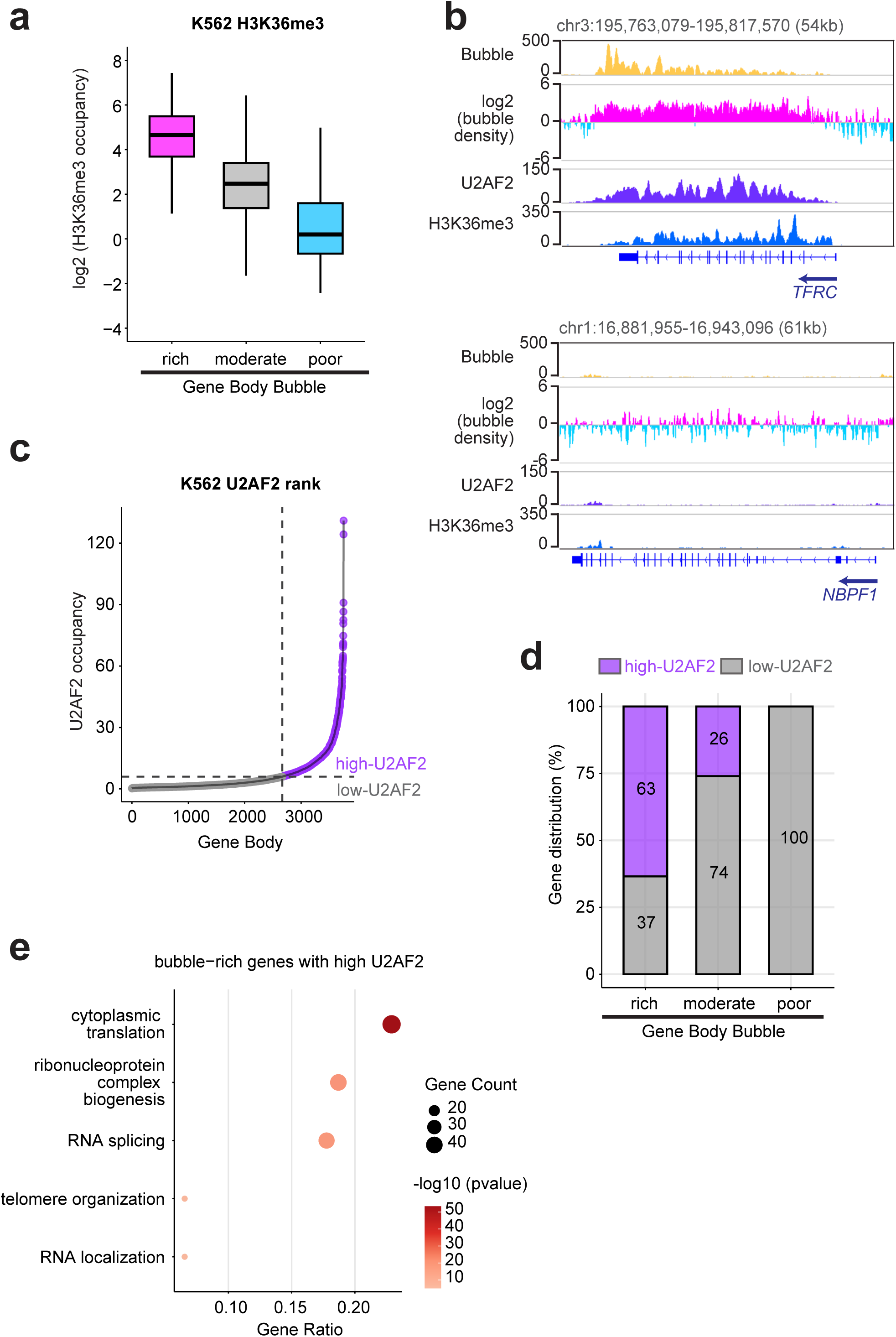
Transcription bubbles are enriched at gene bodies marked by the H3K36me3 mark and the U2AF2 splicing factor. **a,** Coverage-normalized H3K36me3 CnT signals across bubble-rich, -moderate, and -poor gene bodies in K562 cells. H3K36me3 signals in rich (*n* = 342), moderate (*n* = 3,349), and poor (*n* = 63) active gene body bubble groups; medians were 4.66, 2.47, and 0.20, respectively. All pairwise comparisons were significant (two-sided Wilcoxon rank-sum tests, Benjamini–Hochberg-adjusted *p* ≤ 2.77 × 10⁻^17^). Boxes span the 25th–75th percentiles, centre lines show medians, and whiskers extend to the minimum and maximum values within 1.5× the interquartile range; outliers are not shown. **b,** Coverage-normalized signals for Pol II-Ser2p KAS-CnT IP, log₂(Pol II-Ser2p bubble density), U2AF2 CUT&RUN (CnR) and H3K36me3 CnT in K562. **c,** Ranking of 3,754 active genes by U2AF2 CnR signal across gene bodies. **d,** Proportion of genes with high or low U2AF2 signal within bubble-rich, -moderate, and -poor groups. **e,** Top five nonredundant GO terms enriched among bubble-rich genes with high U2AF2 binding. Enrichment was assessed using a one-sided hypergeometric test with Benjamini–Hochberg correction. Dot position indicates gene ratio, dot size indicates gene count, and color indicates −log₁₀(BH-adjusted *P* value).

The H3K36 methyltransferase SETD2^37^ binds elongating Pol II^38,39^, and deposits H3K36me3 to nucleosomes partially unwrapped by the Pol II transcription machinery^40^. One function of the H3K36me3 modification is to recruit the U2AF2 splicing factor that travels with elongating Pol II to chromatin^41^. This chromatin-bound U2AF2 preferentially binds highly expressed genes to enhance co-transcriptional splicing^41^. Comparing the distributions of Pol II-Ser2p bubble density, H3K36me3, and U2AF2 binding reveals that these three features co-occupy many genes in K562 cells. For example, the *TFRC* gene had high Pol II-Ser2p bubble density, H3K36me3, and U2AF2 binding throughout (Fig. 5b). In contrast, the *NBPF1* gene had little Pol II-Ser2p bubble density, H3K36me3, or U2AF2. Genome-wide analysis confirms that U2AF2 occupancy was high in bubble-rich gene bodies (Extended Data Fig.6b). We found that 63% (217 of 342) of bubble-rich and 26% (872 of 3,349) of bubble-moderate gene bodies showed high U2AF2 occupancy, compared to none of the bubble-poor gene bodies (Fig. 5c,d). Gene Ontology analysis on the 217 bubble-rich genes with high U2AF2 found that 18% of these genes are associated with RNA splicing (Fig. 5e).

These features also coincide in HEK293 cells. For example, the *HNRNPU* gene showed higher transcription bubble density, H3K36me3 and U2AF2 occupancy, and nascent RNA levels than that at the *ZDHHC16* gene (Extended Data Fig.6c). Genome-wide, U2AF2 levels were ∼10-fold higher at bubble-rich and 4-fold higher at bubble-moderate gene bodies than at bubble-poor genes (Extended Data Fig.6d). These findings suggest that Pol II-Ser2p generates more transcription bubbles at genes marked by H3K36me3 and bound by U2AF2.

### Transcription factor NPAT is juxtaposed to transcription bubbles at the Histone Locus Body

To identify genes with high transcription bubble density, we ranked the 3,859 active genes by bubble density. Remarkably, 90% (51/57) of active replication-coupled (RC) histone genes ranked within the top 10%, indicating exceptionally high transcription bubble density in K562 cells (Fig. 6a). Similarly, 82% (47/57) of active RC histone genes ranked within the top 10% by PRO-seq signal, indicating exceptionally high nascent transcription in K562 cells (Fig. 6b). High transcription bubble density and nascent transcription levels were also observed at RC histone genes in GM12878 cells (Fig. 6c, d).

**Fig. 6:**
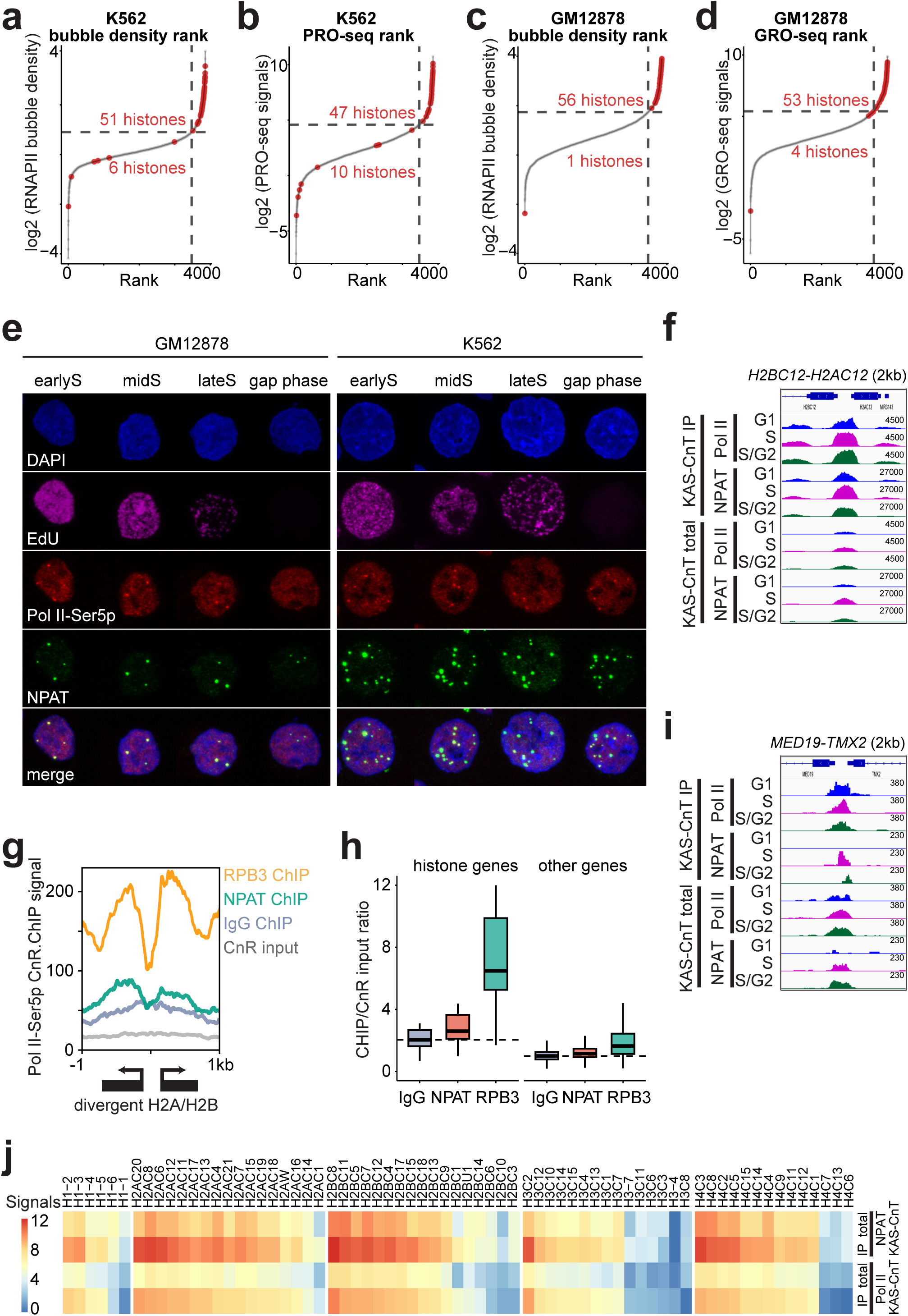
NPAT is juxtaposed to transcription bubbles at the Histone Locus Body. **a, b,** Ranking of 3,859 active genes by log₂(Pol II bubble density), quantified as log₂(IP/total), across the TSS-to-TES region in K562 (**a**) and GM12878 (**c**) cells. Red dots indicate RC histone genes, and gray dots indicate other active genes. **b, d,** Ranking of 3,859 active genes by log₂(PRO-seq signal) in K562 cells (**b**) and log₂(GRO-seq signal) in GM12878 cells (**d**), quantified across the TSS-to-TES region. **e,** Immunostaining of human GM12878 and K562 cells for the HLB factor NPAT (green) and Pol II-Ser5p (red), with EdU detected by click chemistry (magenta). For display, identical minimum and maximum intensity settings were applied to all images within each cell type and channel. At least 300 cells were analyzed. NPAT–Pol II-Ser5p colocalization foci were observed mainly in S-phase GM12878 cells, but throughout interphase in K562 cells. **f,i,** Coverage-normalized signals for Pol II KAS-CnT IP and total at representative bidirectional H2A/H2B gene pairs (**f**) and bidirectional non-histone gene pairs (**i**) in G1, S, and S/G2-phase K562 cells. **g,** Coverage-normalized CUT&RUN.ChIP signals for the Pol II-Ser5p CUT&RUN (CnR) input and its subsequent IgG, NPAT, and RPB3 ChIP fractions across divergent H2A/H2B histone gene pairs. **h,** Ratios of IgG, NPAT, and RPB3 ChIP signals to Pol II-Ser5p CnR input at RC histone and other active gene TSSs. The dashed line marks the median IgG ratio as nonspecific background. Boxplot center lines show medians; IgG, NPAT, and RPB3 ratios were 2.03, 2.59, and 6.48 at histone TSSs and 1.00, 1.15, and 1.63 at other active TSSs, respectively. All pairwise comparisons within each category were significant (two-sided Wilcoxon rank-sum tests, Benjamini–Hochberg-adjusted *P* ≤ 1.05 × 10⁻^3^). Boxes span the 25th–75th percentiles, centre lines show medians, and whiskers extend to the minimum and maximum values within 1.5× the interquartile range; outliers are not shown. **j,** Heat maps showing coverage-normalized Pol II and NPAT KAS-CnT IP and total signals at individual RC histone genes in S-phase K562 cells. Rows were sorted by Pol II KAS-CnT IP signal within each histone type.

Histone locus bodies (HLBs) are nuclear compartments that assemble at RC histone genes and require NPAT for their formation and histone gene activation^42^. We immunostained GM12878 and K562 cells for NPAT and initiating Pol II (Pol II-Ser5p). EdU staining patterns distinguished early, mid, and late S phase (Fig. 6e), while EdU-negative interphase cells were classified as gap phase cells (Extended Data Fig.7a). Approximately 34% of GM12878 and 65% of K562 cells were in S phase (Extended Data Fig.7b). In GM12878 cells, approximately 90% of S-phase and 21% of gap phase cells contained at least one colocalized Pol II-Ser5p and NPAT focus, compared with 75% and 70%, respectively, in K562 cells (Fig. 6e; Extended Data Fig.7c). Pol II KAS-CUT&Tag total signals remained enriched at histone genes across FACS-sorted G1, S, and S/G2 populations of K562 cells, with IP signals exceeding total signals (Fig. 6f; Extended Data Fig.7d). Together, these findings indicate that Pol II localizes to histone genes mainly during S phase in GM12878 cells but remains associated with transcription bubbles at these loci throughout interphase in K562 cells.

To test whether NPAT and Pol II-Ser5p co-occupy histone genes in native chromatin, we performed CUT&RUN.ChIP^43,44^, in which CUT&RUN releases chromatin particles containing one epitope and ChIP detects a second epitope on those particles. Following Pol II-Ser5p CUT&RUN, NPAT ChIP recapitulated the enrichment pattern of the Pol II subunit RPB3 at H2A/H2B gene pairs (Fig. 6g). We calculated ChIP/input ratios, using the IgG ratio to estimate nonspecific background. RPB3 ratios exceeded background at all active gene TSSs, whereas NPAT ratios exceeded background only at RC histone gene TSSs (Fig. 6h). These results support specific NPAT–Pol II-Ser5p interactions at histone genes.

We next profiled NPAT chromatin occupancy and association with transcription bubbles in K562 cells using KAS-CUT&Tag total and IP signals, respectively. NPAT and Pol II IP signals coincided at bidirectional H2A/H2B histone gene pairs and were much stronger than at non-histone bidirectional gene pairs (Fig. 6f,i; Extended Data Fig.7e–g). MACS2 peak calling on S-phase NPAT IP data identified 36 peaks at histone gene clusters on chromosomes 1 and 6. Pol II IP signals were centered on these peaks across G1, S, and S/G2 populations, with both NPAT and Pol II IP signals exceeding their respective total signals (Extended Data Fig.7g). NPAT and Pol II IP signals also showed concordant enrichment across all RC histone genes (Fig. 6j). These results indicate that NPAT is juxtaposed to transcription bubbles at histone genes throughout interphase in K562 cells.

To examine NPAT association with transcription bubbles around HLBs beyond RC histone genes, we analyzed the 36 strongest S-phase NPAT IP peaks near HLBs that did not overlap RC histone genes, using the 36 strongest peaks outside HLBs as controls. Pol II KAS-CUT&Tag IP signals were centered on HLB-proximal NPAT peaks (Extended Data Fig.8a), supporting NPAT association with transcription bubbles at non-histone sites. After excluding three control peaks containing tandem AGGGTT repeats, Pol II IP signals were also centered on the remaining 33 NPAT peaks (Extended Data Fig.8b). Of these, 13 were within 0.5 Mb of the major histone gene cluster *HIST1*, and four others were elsewhere on chromosome 6p (Extended Data Fig.8b). Thus, 17 of 33 peaks (∼52%) were on chromosome 6p, which comprises only ∼2% of the genome. This enrichment suggests that HLB proximity may favor NPAT association with transcription bubbles in K562 cells. Together, these results show that KAS-CUT&Tag detects NPAT and Pol II co-occupying transcription bubbles at HLBs in K562 cells.

## Discussion

KAS-CUT&Tag is a rapid, low-input, and easily standardized method, enabling direct and efficient mapping of transcription bubbles. While KAS-seq maps ssDNA and can be used to infer transcription bubble locations, its signals are not specific to bubbles formed by Pol II and may also arise from DNA damage, R-loops, non-canonical structures, or DNA replication^17^. KAS-ATAC-seq increases specificity by combining ssDNA labeling with Tn5 tagmentation but cannot resolve protein-specific interactions^21,22^. RNA-based methods, including RNA-seq^45^, GRO-seq^46,47^, PRO-seq^48^, NET-seq^49^, mNET-seq^50^, SLAM-seq^51^, and TimeLapse-seq^52^, indirectly infer Pol II dynamics via steady-state or nascent transcripts, often requiring high cell input and showing limited sensitivity to low-abundance RNAs.

Using an antibody raised against the unmodified Pol II CTD, KAS-CUT&Tag provides an unbiased and direct readout of transcription bubble dynamics across all cellular contexts. We found that inhibition of pause release increased bubble density at promoters, while blocking the reactivation of backtracked Pol II led to reduced bubble density at promoters and post-transcription end sites, demonstrating the sensitivity of Pol II KAS-CUT&Tag in detecting general transcriptional dynamics. Traditionally, extracellular signals are thought to regulate transcription via intracellular intermediates that activate transcription factors, which then regulate Pol II activity at target genes. However, recent studies have shown that nearly one-quarter of the ∼500 human tyrosine kinases can directly phosphorylate the Pol II CTD at the Y1, T4, or S7 residues, with each kinase targeting distinct promoter subsets, likely via specific transcription factors interactions^53^. Detection of these modifications has been limited by poor antibody performance, due to the close proximity of the Y1, T4, and S7 residues to other phosphorylated positions. Pol II KAS-CUT&Tag overcomes this limitation by directly profiling transcription bubbles without relying on CTD phosphorylation, making it a powerful tool to investigate the effects of kinase and transcription factors perturbations. KAS-CUT&Tag is general, and should be adaptable to other eukaryotic and prokaryotic RNA polymerases. Additionally, KAS-CUT&Tag may be extended to specifically profile single-stranded regions during replication, recombination and repair with suitable antibodies.

KAS-CUT&Tag also enables detection of TFs and regulatory proteins juxtaposed to transcription bubbles through their association with Pol II. Because Pol II generates transcription bubbles, the precisely coincident NPAT and Pol II KAS-CUT&Tag signals indicate NPAT–Pol II co-occupancy, which was independently validated by Pol II-Ser5p CUT&RUN.ChIP. NPAT and Pol II co-occupy not only histone genes but also non-genic sites within and near the HLB, where their signals were approximately two orders of magnitude above background. NPAT is a largely disordered 1427 aa protein^54^, and the Pol II CTD is also intrinsically disordered. Because weak heterotypic interactions between IDRs can stabilize TF binding to chromatin^55^, we infer that such interactions may mediate NPAT–Pol II juxtaposition at transcription bubbles.

We have shown that, in GM12878 cells, Pol II accumulates at HLBs during S phase and is depleted in G1 and G2 phases, whereas this S-phase specificity is lost in K562 cells (Fig. 6d). Furthermore, in K562 cells, Pol II remains engaged at transcription bubbles throughout interphase, and NPAT likewise remains juxtaposed to transcription bubbles at histone gene promoters (Fig. 7e). This persistent Pol II–NPAT association at transcription bubbles may facilitate G1-to-S transitions through CDK2-mediated histone gene activation in K562 cells. Consistent with this model, only 11% of K562 cells were in G1 and 65% were in S phase, whereas 39% of GM12878 cells were in G1 and 34% were in S phase. Given that Pol II upregulation of RC histone genes^56^ and shortened total cell-cycle duration^57^ are CDK2-dependent cancer-associated features, one possible explanation is that Pol II initiation at histone genes is normally restricted to S phase to tightly couple histone production to DNA replication, whereas persistent Pol II–NPAT association at histone gene transcription bubbles in cancer cells may drive accelerated cell-cycle progression.

## Acknowledgements

We thank Jorja Henikoff for help with processing and analysis of sequencing data, Christine Codomo for assistance with sequencing library pooling, Terri Bryson and Doris Xu for technical support, and Ronald Paranal for assistance with figure modifications. We also thank members of the Henikoff laboratory for their valuable feedback and discussions throughout the course of this work, and Evgeny Nudler (New York University) for providing the TFIIS KO cell line. We thank Lena Schroeder and Hoku West-Foyle at the Fred Hutch Cellular Imaging Facility for assistance with the Leica Stellaris 8 scanning confocal microscope, and Lena Schroeder for assistance with fluorescence image analysis.

## Contributions

K.A. and S.H. supervised the project. W.W. designed and implemented the experiments. W.W. conducted the computational analyses. W.W. wrote the paper with input from K.A., and S.H. J.G. performed Fluorescence-Activated Cell Sorting of K562 cells.

## Methods

### Human cell culture

Human K562 (ATCC CCL-243), HEK293T (ATCC CRL-11268), and GM12878 (Coriell Institute) cells were cultured at 37 °C with 5% CO_2_ according to supplier recommendations. K562 cells were maintained in IMDM (ATCC, 30-2005) with 10% FBS (HyClone, SH30070.03); HEK293T cells in high-glucose DMEM (Gibco, 10566016) with 10% FBS and 1× Antibiotic-Antimycotic (Gibco, 15240062); and GM12878 cells in RPMI-1640 containing 2 mM L-glutamine (Gibco, 11875093) and 15% FBS. For transcription inhibition, K562 cells were treated for 1 h with triptolide (10 μM; Selleckchem, S3604), flavopiridol hydrochloride hydrate (1 μM; Sigma-Aldrich, FL3055), or actinomycin D (5 μg ml⁻^1^; Sigma-Aldrich, A9415), with DMSO (1:1,000, v/v) as the vehicle control, and then processed for KAS-CUT&Tag.

### Antibodies

Primary antibodies were H3K4me3 (Rabbit monoclonal, Epicypher, 13-0041); H3K27me3 (Rabbit monoclonal, Cell Signaling Technology, 9733 BC); H3K36me3 (Rabbit monoclonal, Epicypher, 13-0058); Pol II (Mouse monoclonal, Sigma-Aldrich, 05-952-I); Pol II-Ser2p (Rabbit monoclonal, Cell Signaling Technology, 13499S); Pol II-Ser5p (Rabbit monoclonal, Cell Signaling Technology, 13523S); U2AF2 (Rabbit polyclonal, Abcam, ab37530); NPAT (Rabbit polyclonal, Invitrogen, PA565419) and NPAT (Mouse monoclonal, Abcam, ab307837); RPB3 (Rabbit polyclonal, Bethyl Laboratories, A303-771A). Secondary antibodies were Guinea Pig anti-Rabbit IgG H&L (Antibodies Online, ABIN101961); Rabbit Anti-Mouse IgG H&L (Abcam, ab46540); Alexa Fluor® 488 AffiniPure® Fab Fragment Goat Anti-Mouse IgG (Jackson ImmunoResearch, 115-547-003) and Rhodamine Red™-X (RRX) AffiniPure® Fab Fragment Goat Anti-Rabbit IgG (Jackson ImmunoResearch, 111-297-003).

### Fluorescence-Activated Cell Sorting

To purify G1, S, G2 population of K562 cells, 10 million cells were resuspended in ice-cold harvest buffer (1x PBS with 2 mM EDTA, 2% FBS, 1 mM Spermidine, 0.05% Triton-X-100) and briefly fixed with 0.1% PFA for 2 minutes. Fixation was quenched with 2.5 M glycine for 5 minutes, then cells were spun down and resuspended in harvest buffer with 2.5 μM DRAQ5 (Thermo Scientific, 62251) to stain DNA. After at least 10 minutes of incubation, three cell cycle fractions were isolated on a BD Discover S8 imaging flow cytometer based on DRAQ5 intensity.

### Immunofluorescence staining

K562 and GM12878 cells were processed for immunofluorescence staining as follows. For EdU labeling, cells were incubated with 10 μM EdU in growth medium for 15 min before harvest. 500,000 cells were pelleted and resuspended in 500 μL of 4% formaldehyde (4% formaldehyde in 1× PBS; Thermo Scientific, 28906) and fixed on ice for 5 min. Fixation was quenched by addition of 25 μL of 2.5 M glycine, followed by centrifugation. Cells were then resuspended in 1 mL of PBST (1× PBS, 0.1% Triton X-100), pelleted, and resuspended again in 1 mL of 1× PBS. For slide preparation, 50 μL of cell suspension (approximately 50,000 cells) was deposited onto each glass slide by cytospin and immediately washed in 1× PBS in a Coplin jar. Cells were then blocked for 30 min at room temperature in 100 μL of wash blocking buffer (20 mM HEPES, pH 7.5, 150 mM NaCl, 0.5 mM spermidine, 0.1% Triton X-100, one EDTA-free protease inhibitor tablet per 50 mL (Sigma-Aldrich, 05056489001), 0.5% BSA (Sigma-Aldrich, A7906), and 0.5% casein (Sigma-Aldrich, C7078)). Rabbit anti-Pol II-Ser5p (Cell Signaling Technology, 13523S) and mouse anti-NPAT (Abcam, ab307837) primary antibodies were diluted 1:200 in antibody blocking buffer (wash blocking buffer supplemented with 2 mM EDTA) and pre-incubated for 30 min at 4°C. After blocking, 20 μL of the primary antibody dilution was added to each slide, and slides were incubated overnight at 4°C in a humid chamber. The following day, slides were washed in 1× PBS in a Coplin jar and blocked again for 30 min at room temperature in the same wash blocking buffer. Secondary antibodies (anti-rabbit Rhodamine and anti-mouse Fab-Alexa Fluor 488) were diluted 1:500 in wash blocking buffer and pre-incubated for 30 min at 4°C. 20 μL of the secondary antibody dilution was then added to each slide, followed by incubation for 1 h at 4°C. After secondary antibody incubation, slides were washed in 1× PBS and subjected to EdU detection using the Invitrogen Click-iT Plus EdU Cell Proliferation Kit with Alexa Fluor 647 dye (Fisher Scientific, C10640), according to the manufacturer’s instructions. The click reaction was carried out for 1 h at room temperature protected from light, followed by two washes in 1× PBS. Nuclear DNA was stained with 1× DAPI for 5 min at room temperature. Slides were then washed in 1× PBS, mounted in 80% glycerol, coverslipped, and sealed with nail polish before imaging.

### Fluorescence image acquisition and analysis

Images were acquired using a Leica Stellaris 8 scanning confocal microscope equipped with a 63×/1.4 NA oil HC PL APO CS2 objective. Z-stacks were collected from K562 and GM12878 cells. DAPI, Alexa Fluor 488, Rhodamine, and Alexa Fluor 647 were excited with 405 nm, 495 nm, 571 nm, and 649 nm lasers, respectively, and detected at 429–502 nm, 503–567 nm, 579–646 nm, and 654–783 nm, respectively. Images were analyzed using Imaris v11.0.1 for cell segmentation and detection of Pol II-Ser5p and NPAT foci, and the number of colocalized foci per cell was quantified using Imaris Cell. Colocalized foci were defined as Pol II-Ser5p foci with NPAT mean intensity greater than 30. EdU staining patterns were used to classify cells as early, mid, or late S phase, while EdU-negative interphase cells were classified as gap phase cells. Pseudocolored maximum-intensity projections were generated from z-stacks. For display, identical minimum and maximum intensity settings were applied to all images within each cell type and channel, and images were composited in Fiji. Images exported from Fiji were assembled into the final figures using Adobe Illustrator.

### CUT&RUN

CUT&RUN was carried out following a previously described protocol^58^, with several modifications. A total of 600,000 cells were collected by centrifugation and resuspended in wash buffer (20 mM HEPES pH 7.5, 150 mM NaCl, 0.5 mM spermidine, and one EDTA-free protease inhibitor tablet per 50mL (Sigma-Aldrich, 05056489001)). The cell suspension was then incubated with activated concanavalin A (ConA)-coated magnetic beads (5 μl per sample; Bangs Laboratories, BP531) for 10 minutes at room temperature to allow bead binding. Following this, the bead-bound cells were aliquoted into individual PCR tubes, the supernatant was removed, and the beads were resuspended in 50 μl of antibody buffer (wash buffer containing 0.01% Digitonin and 2 mM EDTA). Primary antibody against U2AF2, H3K4me3 or H3K27me3 were added at a 1:20 dilution, and samples were incubated overnight at 4 °C. After antibody binding, the samples were washed twice with dig wash buffer (wash buffer + 0.01% Digitonin). They were then incubated for 10 minutes at room temperature in 50 μl of dig wash buffer with Protein A–MNase (pA-MN, prepared in-house) at a 1:130 dilution, followed by four additional washes with 200 μl of dig wash buffer. To initiate MNase digestion, the cell-bead slurry was resuspended in 50 μl of dig wash buffer containing 2 mM CaCl_2_ and incubated at 4 °C for 2 hours. The digestion reaction was stopped by adding 33 μl of stop buffer (340 mM NaCl, 20 mM EDTA, 4 mM EGTA, 50 μg/ml glycogen (Sigma-Aldrich, 10930193001), 50 μg/ml RNase A (Thermo Fisher Scientific, EN0531), and 2 pg/ml *E. coli* spike-in DNA), followed by a 10-minute incubation at 37 °C to release DNA fragments. The supernatant was collected, and DNA was purified using the CUTANA™ DNA Purification Kit (EpiCypher, 14-0050) according to the manufacturer’s instructions. The resulting DNA was then used for library preparation as described previously^58^.

### CUT&RUN.CHIP

CUT&RUN.ChIP was performed as described previously^43,44^, with modifications. For each experiment, 2.5 × 10^6^ K562 cells were bound to 50 µl of ConA-coated magnetic beads. Incubation with the Pol II-Ser5p primary antibody (Cell Signaling Technology, 13523S), washes and pA–MN binding were performed as described above for CUT&RUN, using five times the corresponding buffer volumes in 1.5-ml low-binding flip-cap tubes. MNase digestion was initiated by resuspending the ConA beads in 500 µl of dig-wash buffer containing 2 mM CaCl₂ and incubating for 2 h at 4 °C. The reaction was stopped by directly adding 500 µl of stop buffer supplemented with Pol II CTD repeat YSPTSPS phospho-Ser5 peptide (Abcam, ab18488) to a final concentration of 20 µg/ml. Samples were incubated for 20 min at 37 °C and centrifuged for 5 min at 16,000g and 4 °C. The 1 ml of supernatant containing released Pol II-Ser5p-bound native chromatin fragments was collected on a magnetic stand and divided into five 200-µl aliquots. One aliquot was retained at 4 °C as the Pol II-Ser5p CUT&RUN input, and the remaining aliquots were used for ChIP. Rabbit monoclonal IgG isotype-control (Abcam, ab172730), anti-NPAT (Invitrogen, PA565419) or anti-RPB3 (Bethyl Laboratories, A303-771A) antibody was added to the corresponding ChIP aliquot at a final dilution of 1:100, and samples were incubated overnight at 4 °C. Protein A Dynabeads (Invitrogen, 10002D) were equilibrated in Tween wash buffer consisting of wash buffer supplemented with 0.05% Tween-20 (Sigma, P7949). 20 µl of Dynabeads was added to each ChIP sample, followed by incubation for 30 min at 4 °C. Beads were washed once with 500 µl of Tween wash buffer and resuspended in 5 µl of release solution containing 10 mM TAPS, 0.1% SDS and Thermolabile Proteinase K at a 1:10 dilution (New England Biolabs, P8111S). Samples were incubated in a thermocycler with a heated lid for 1 h at 37 °C and then for 1 h at 58 °C. SDS was quenched by adding 4 µl of 1.5% Triton X-100 and and 24 μl of ddH_2_O. DNA from the input and ChIP samples was purified using the DNA Clean & Concentrator-5 Kit (Zymo Research, D4014), eluted in 10 µl of elution buffer according to the manufacturer’s instructions and used for library preparation as described previously^58^. A step-by-step protocol used in this study, modified from the published protocol^59^, is available through the Gene Expression Omnibus under accession number GSE317877.

### CUT&Tag

We performed CUT&Tag based on the CUT&Tag-direct protocol^60,61^ with slight modifications. To begin, 30,000 cells were pelleted and gently resuspended in wash buffer (20 mM HEPES pH 7.5, 150 mM NaCl, 0.5 mM spermidine, 0.05% Triton X-100, one EDTA-free protease inhibitor tablet per 50mL (Sigma-Aldrich, 05056489001)). ConA-coated magnetic beads (5 μl per sample; Bangs Laboratories, BP531) were pre-washed and activated in beads binding buffer (20 mM HEPES pH 7.9, 10 mM KCl, 1 mM CaCl_2_, 1 mM MnCl_2_), then added to the cell suspension and incubated at room temperature for 10 minutes. Bead-bound cells were transferred to 20 μl of antibody binding buffer (wash buffer + 2 mM EDTA), split into 0.5 ml tubes, and incubated overnight at 4 °C with primary antibody against H3K36me3 (Epicypher, 13-0058), Pol II (Sigma-Aldrich, 05-952-I), or Pol II-Ser5p (Cell Signaling Technology, 13523S) at a final dilution of 1:10. Following incubation, unbound antibody was removed by washing with 200 μl of wash buffer, and bead-bound cells were subsequently incubated in 30 μl of wash buffer containing secondary antibody (1:15 dilution) for 1 hour at 4 °C. After another wash with 200 μl of wash buffer, beads were resuspended in 20 μl of 300-wash buffer (wash buffer + 150 mM NaCl) containing Protein A/G-Tn5 (1:10 dilution; EpiCypher, 15-1117) and incubated for 1 hour at 4 °C. Unbound pAG-Tn5 was removed by washing using 200 μl of 300-wash buffer, and tagmentation was performed by resuspending beads in 50 μl of 300-tagmentation buffer (300-wash buffer containing 10 mM MgCl_2_) and incubating at 37 °C for 1 hour. Following tagmentation, beads were washed with 100 μl of TAPS buffer (10 mM TAPS, 0.2 mM EDTA) and resuspended in 5 μl of release solution (10 mM TAPS, 0.1% SDS, and 1:10 Thermolabile Proteinase K; New England Biolabs, P8111S). Samples were incubated in a thermocycler with a heated lid for 1 h at 37 °C, followed by 1 h at 58 °C, to release the tagmented DNA fragments. SDS was quenched by adding 4 µl of 1.5% Triton X-100 and 24 µl of ddH₂O. For PCR amplification, 33 µl of the released DNA was combined with 22 µl of PCR mix containing 2 µl each of 10 µM barcoded i5 and i7 primers (IDT), 10 µl of 5× HiFi buffer, 1.5 µl of 10 mM dNTPs, 1 µl of KAPA HiFi polymerase and 5.5 µl of H₂O (Roche, 07958846001), for a final reaction volume of 55 µl. PCR cycling conditions were: 58 °C for 5 min; 72 °C for 5 min; 98 °C for 30 s; followed by 12 cycles of 98 °C for 10 s and 60 °C for 10 s; then 72 °C for 1 min and hold at 12 °C. Libraries were purified using HighPrep paramagnetic beads (MagBio, AC-60500) at a 1.3:1 bead-to-sample volume ratio.

### KAS-CUT&Tag

A total of 30,000 cells were pelleted and gently resuspended in antibody binding buffer (20 mM HEPES pH 7.5, 150 mM NaCl, 0.5 mM spermidine, 0.05% Triton X-100, 2 mM EDTA, and EDTA-free protease inhibitor tablet). Separately, 5 μl of ConA-coated magnetic beads (Bangs Laboratories, BP531) were washed and activated in beads binding buffer (20 mM HEPES pH 7.9, 10 mM KCl, 1 mM CaCl_2_, 1 mM MnCl_2_), then added to the cell suspension and incubated for 10 minutes at room temperature. After binding, bead-bound cells were resuspended in 200 µl of fresh antibody binding buffer in 0.5 ml low-binding tubes. The buffer was removed, and the cells were incubated in 100 µl of 1× PBS containing 10 mM N_3_-kethoxal (ApexBio, A8793) for 10 min at 37 °C. Cells were then washed twice in 200 μl of DPBS and fixed in 20 μl of 0.1% formaldehyde (Thermo Scientific, 28906) for 1 minute. Fixation was omitted when tagmentation was subsequently performed in 300-tagmentation buffer (300-wash buffer containing 10 mM MgCl₂). Following fixation, cells were rinsed in 100 μl of antibody binding buffer and incubated overnight at 4 °C in 20 μl of antibody binding buffer with primary antibody against Pol II (Sigma-Aldrich, 05-952-I), Pol II-Ser5p (Cell Signaling Technology, 13523S), Pol II-Ser2p (Cell Signaling Technology, 13499S) or NPAT (Invitrogen, PA565419) at a final dilution of 1:10. The next day, cells were washed to remove unbound antibody by 200 μl of wash buffer (20 mM HEPES pH 7.5, 150 mM NaCl, 0.5 mM spermidine, 0.05% Triton X-100, one EDTA-free protease inhibitor tablet per 50mL (Sigma-Aldrich, 05056489001)) and incubated in 30 μl of wash buffer with secondary antibody (1:15 dilution) for 1 h at 4 °C. After washing by 200 μl of wash buffer, beads were resuspended in 20 μl of 300-wash buffer (wash buffer + 50 mM NaCl) containing pAG-Tn5 transposase (1:10 dilution; EpiCypher, 15-1117) and incubated at 4 °C for 1 h. Excess pAG–Tn5 was removed by washing twice with 200 µl of 300-wash buffer. Samples fixed with 0.1% formaldehyde were tagmented in 50 µl of CUTAC–DMF tagmentation buffer (20% N,N-dimethylformamide (Sigma-Aldrich, D8654), 10 mM TAPS PH8.5, 0.05% Triton X-100, 5 mM MgCl_2_). Unfixed samples were tagmented in 50 µl of 300-tagmentation buffer. Tagmentation was performed for 1 h at 37 °C. Following tagmentation, beads were washed with 200 μl of TAPS buffer (10 mM TAPS, 0.2 mM EDTA) and resuspended in 5 μl of release solution (10 mM TAPS, 0.1% SDS, and 1:10 Thermolabile Proteinase K (New England Biolabs, P8111S). Samples were incubated at 37 °C for 1 h, followed by 58 °C for 1 h in a thermocycler with a heated lid. SDS was quenched by adding 4 μl of 1.5% Triton X-100 and 24 μl of ddH_2_O. DNA was purified using the DNA Clean & Concentrator-5 Kit (Zymo Research, D4014) and eluted in 10 μl of 25 mM K₃BO₃ (pH 7.0). For biotinylation, the 10 μl of DNA eluate was mixed with 34.5 μl of 25 mM K₃BO₃ (pH 7.0), 5 μl of 10× PBS, and 2.5 μl of 20 mM DBCO-PEG₄-Biotin (Sigma, 760749), and incubated at 37 °C for 1.5 hr. Biotinylated DNA was purified again with the Zymo kit and eluted in 55 μl of 25 mM K₃BO₃ (pH 7.0). A 5 μl aliquot was retained as the KAS-CUT&Tag total sample, and the remaining 50 μl was used to generate the KAS-CUT&Tag IP sample by streptavidin enrichment. For each IP sample, 5 μl of Dynabeads MyOne Streptavidin T1 (Thermo Fisher, 65602) or C1 (Thermo Fisher, 65001) were washed with 1× B&W buffer and resuspended in 50 μl of 2× B&W buffer (10 mM Tris-HCl pH 7.4, 1 mM EDTA, 2 M NaCl, 0.1% Tween-20). The 50 μl of washed bead suspension was combined with 50 µl of biotinylated DNA and incubated for 15 min at room temperature. Beads were then washed five times with 100 μl of 1× B&W buffer and resuspended in 28 μl of nuclease-free water for IP library amplification. The 5µl of total sample was brought to 28 µl with 23 µl of nuclease-free water. For PCR amplification, each 28 µl of total or IP sample was combined with 22 µl of PCR mix containing 2 µl each of 10 µM barcoded i5 and i7 primers, 10 µl of 5× HiFi buffer, 1.5 µl of 10 mM dNTPs, 1 µl of KAPA HiFi polymerase and 5.5 µl of H₂O (Roche, 07958846001), for a final reaction volume of 50 µl. PCR cycling conditions were: 58 °C for 5 min, 72 °C for 5 min, 95 °C for 10 min; followed by 12 cycles of 98 °C for 10 s and 60 °C for 10 s; then 72 °C for 1 min, ending with a 12 °C hold. Final libraries were purified using HighPrep paramagnetic beads (MagBio, AC-60500) at a 1.3:1 bead-to-sample ratio. A step-by-step protocol for KAS-CUT&Tag is available from Protocols.io: dx.doi.org/10.17504/protocols.io.e6nvwn82dvmk/v1

### DNA sequencing and data processing

Library size distributions and molar concentrations were assessed using an Agilent 4200 TapeStation. Libraries were pooled to ensure equal representation and adjusted to the final concentration recommended by the manufacturer. Paired-end 50 × 50 bp sequencing was performed on an Illumina NovaSeq X Plus platform by the Fred Hutchinson Cancer Center Genomics Shared Resources. Sequencing data were processed as described. (https://www.protocols.io/view/cut-amp-tag-data-processing-and-analysis-tutorial-e6nvw93x7gmk/v1). For processing sequencing data, adapter sequences were trimmed from 50 bp paired-end reads using Cutadapt 4.4^62^ with the specified parameters “-j 8 --nextseq-trim 20 - m 20 -a AGATCGGAAGAGCACACGTCTGAACTCCAGTCA -A AGATCGGAAGAGCGTCGTGTAGGGAAAGAGTGT -Z”

We used Bowtie2 2.5.1^63^ to map the paired-end 50 bp reads to the hg19 human genome reference sequence from UCSC with parameters “--very-sensitive-local --soft-clipped-unmapped-tlen -- dovetail --no-mixed --no-discordant -q --phred33 -I 10 -X 1000”. MACS2 peak calling for NPAT KAS-CnT data was performed using macs2 callpeak -t $bed -f BEDPE -g hs --keep-dup all -p 1e- 10 -n $sname.

### Data analysis and visualization

BedGraph files were generated using BEDTools^64^ genomecov, and converted to bigWig format with bedGraphToBigWig. The bigWig files were coverage-normalized by scaling read counts at each base pair to the total size of the reference genome (hg19; 3,095,693,983 bp), such that a uniformly distributed signal would yield a value of 1 across all positions. This normalization enables the identification of regions with enriched protein epitope signals on DNA. Genomic bigWig tracks were visualized using the Integrated Genome Viewer. Heatmaps were produced using deepTools (version 3.5.1)^65^ with the computeMatrix and plotHeatmap functions. Published hg19-aligned bigWig files for K562 H3K36me3 CUT&Tag (GSM8343606 and GSM8343607) and U2AF2 CUT&RUN (GSM8339598–GSM8339602) were downloaded from GEO^41^. K562 H3K9me3 CUT&Tag (GSM4835704 and GSM4842201)^60^, H3K27ac CUT&Tag (GSM3536514)^66^, PRO-seq (GSM1480327)^23^, HEK293T PRO-seq (GSM6735112–GSM6735115)^67^, K562 KAS-seq (GSM6219605)^68^ and GM12878 GRO-seq (GSM1480326)^23^ datasets were downloaded from GEO and reprocessed to generate hg19-aligned bigWig files. K562 rRNA-depleted total RNA-seq data were obtained from ENCODE (ENCFF782PCD)^69^. HEK293T R-loop mapping datasets, including MapR (GSM3406747, GSM3406749 and GSM3406753)^24^, R-loop CUT&RUN (GSM3406755 and GSM3406759)^24^ and R-loop CUT&Tag (GSM4730182 and GSM4730198)^25^, were downloaded, mapped to hg19 and merged across replicates. Paired-end FASTQ files for standard ATAC-seq (SRR10319911, SRR10319912)^20^ and N_3_-kethoxal-treated ATAC-seq (SRR28028139^22^, SRR28028140^22^, SRR28764289^21^) in HEK293T cells were downloaded from the SRA and aligned to the hg19 human reference genome using Bowtie2 (v2.5.1)^63^. SAM files were converted to BAM files using SAMtools^70^, and fragment BED files were generated using BEDTools (v2.30.0)^64^. V-plots show fragment length as a function of the distance between each fragment midpoint and the TSSs of 12,397 RefSeq-annotated protein-coding genes. Twelve bidirectional non-histone gene pairs were obtained from Additional File 1 of Wang et al^71^.

### Quantification and statistical analysis

All experiments were performed with at least two biological replicates. Distributional differences between WT and TFIIS KO cells and between each transcription inhibitor and control were assessed using two-sided Kolmogorov–Smirnov tests with Holm correction; adjusted *P* < 0.001 was considered significant. Pairwise comparisons among bubble groups and among ChIP/input ratios were assessed using two-sided Wilcoxon rank-sum tests with Benjamini–Hochberg correction; adjusted *P* < 0.01 was considered significant. GO-term enrichment *P* values for bubble-rich genes with high U2AF2 binding were Benjamini–Hochberg adjusted and reported as −log_10_(adjusted *P*).

## Data availability

All primary sequence data and interpreted track files for sequence data generated in this study have been deposited at the Gene Expression Omnibus: GSE317877

---

**Extended Data Fig. 1.**
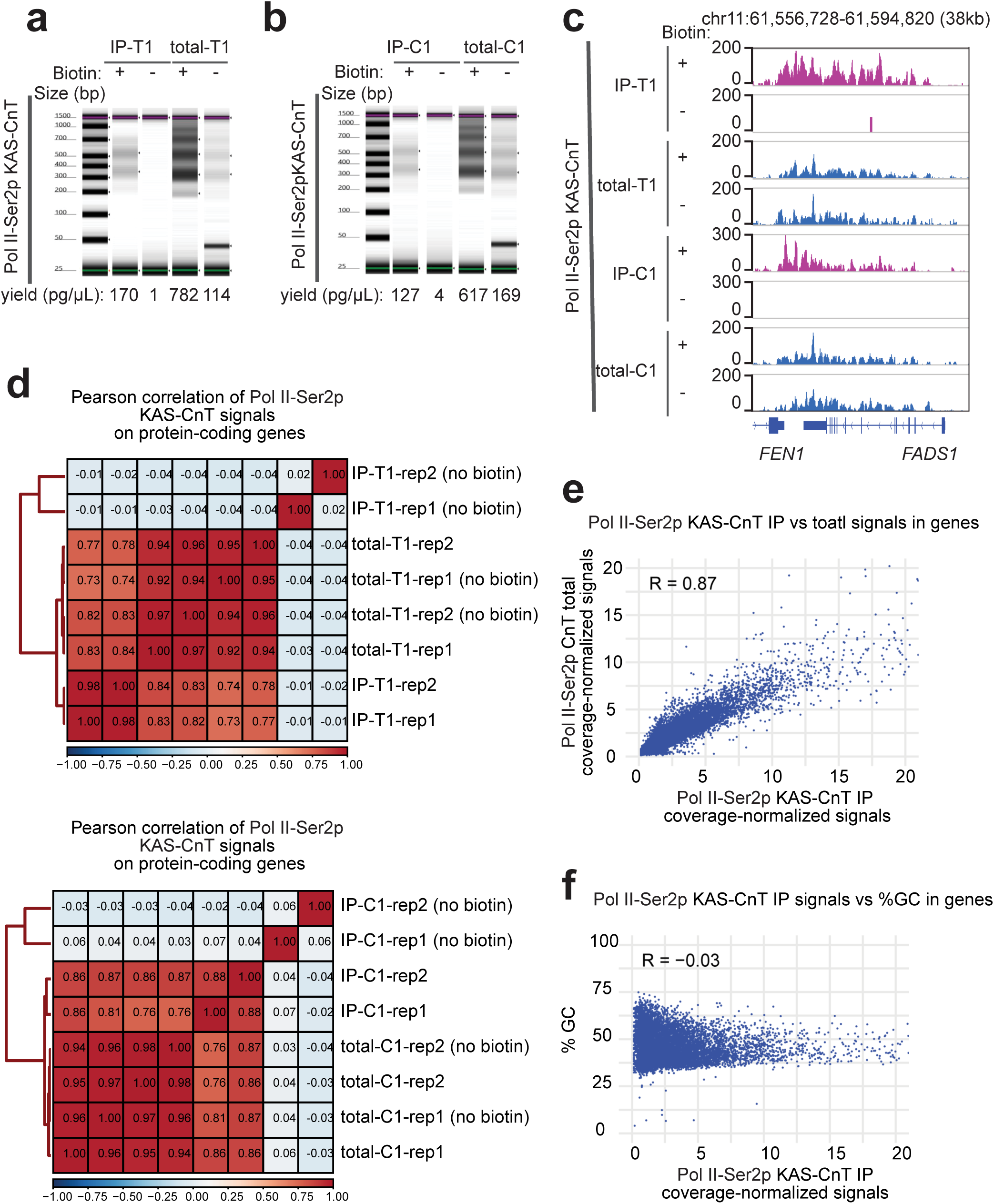
Specificity and reproducibility of KAS-CUT&Tag. a, TapeStation profile of Pol II-Ser2p KAS-CnT IP and total libraries prepared with or without biotin click chemistry in K562 cells using streptavidin T1 beads. b, As in (a), but using streptavidin C1 beads. IP-T1 (a) and IP-C1 (b) denote IP libraries enriched using streptavidin T1 and C1 beads, respectively; total-T1 and total-C1 denote the corresponding total controls. c, Coverage-normalized KAS-CnT IP and total signals from libraries prepared with or without biotin click chemistry in K562 cells. d, Pearson correlations of coverage-normalized Pol II-Ser2p KAS-CnT signals across protein-coding genes. e, Scatter plot comparing coverage-normalized Pol II-Ser2p KAS-CnT IP and total signals across protein-coding genes in K562 cells. Each dot represents one gene. f, Scatter plot showing no relationship between Pol II-Ser2p KAS-CnT IP signal and GC content (%GC) across protein-coding genes in K562 cells. Each dot represents one gene.

**Extended Data Fig. 2.**
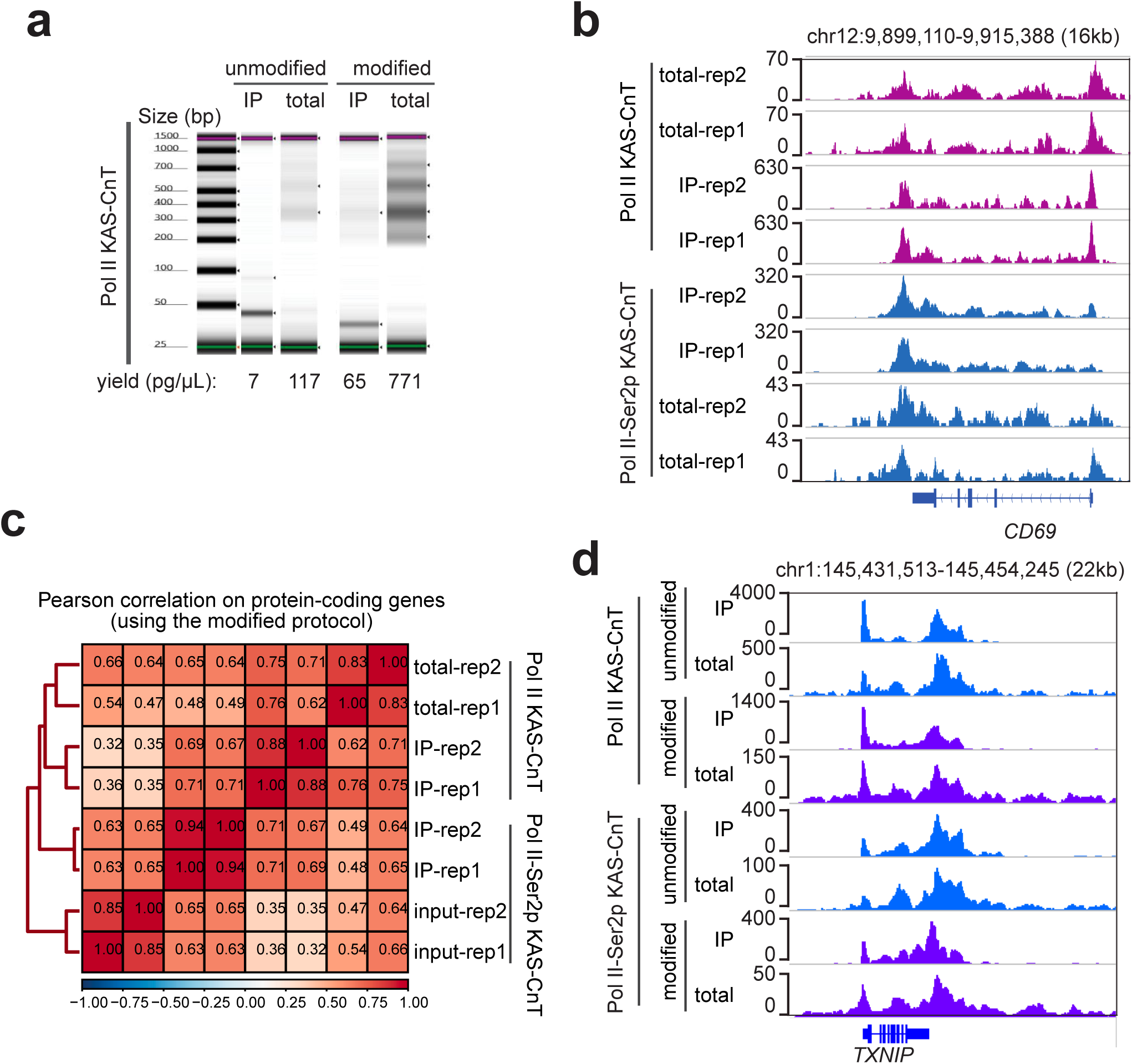
Light crosslinking and CUTAC tagmentation increase KAS-CUT&Tag library yield while maintaining high reproducibility. a, TapeStation profiles of Pol II KAS-CnT IP and total libraries prepared from K562 cells using the modified or unmodified protocol. The modified protocol includes light crosslinking with 0.1% formaldehyde for 1 min after N_3_-kethoxal treatment, followed by tagmentation under low-salt CUTAC conditions. The unmodified protocol omits crosslinking and uses 300 mM NaCl during tagmentation. b, Coverage-normalized Pol II and Pol II-Ser2p KAS-CnT IP and total signals in K562 cells generated using the modified protocol. c, Pearson correlations of coverage-normalized Pol II and Pol II-Ser2p signals across protein-coding genes between replicates generated using the modified protocol. d, Coverage-normalized Pol II and Pol II-Ser2p KAS-CnT IP and total signals in K562 cells generated using the modified or unmodified protocol.

**Extended Data Fig. 3.**
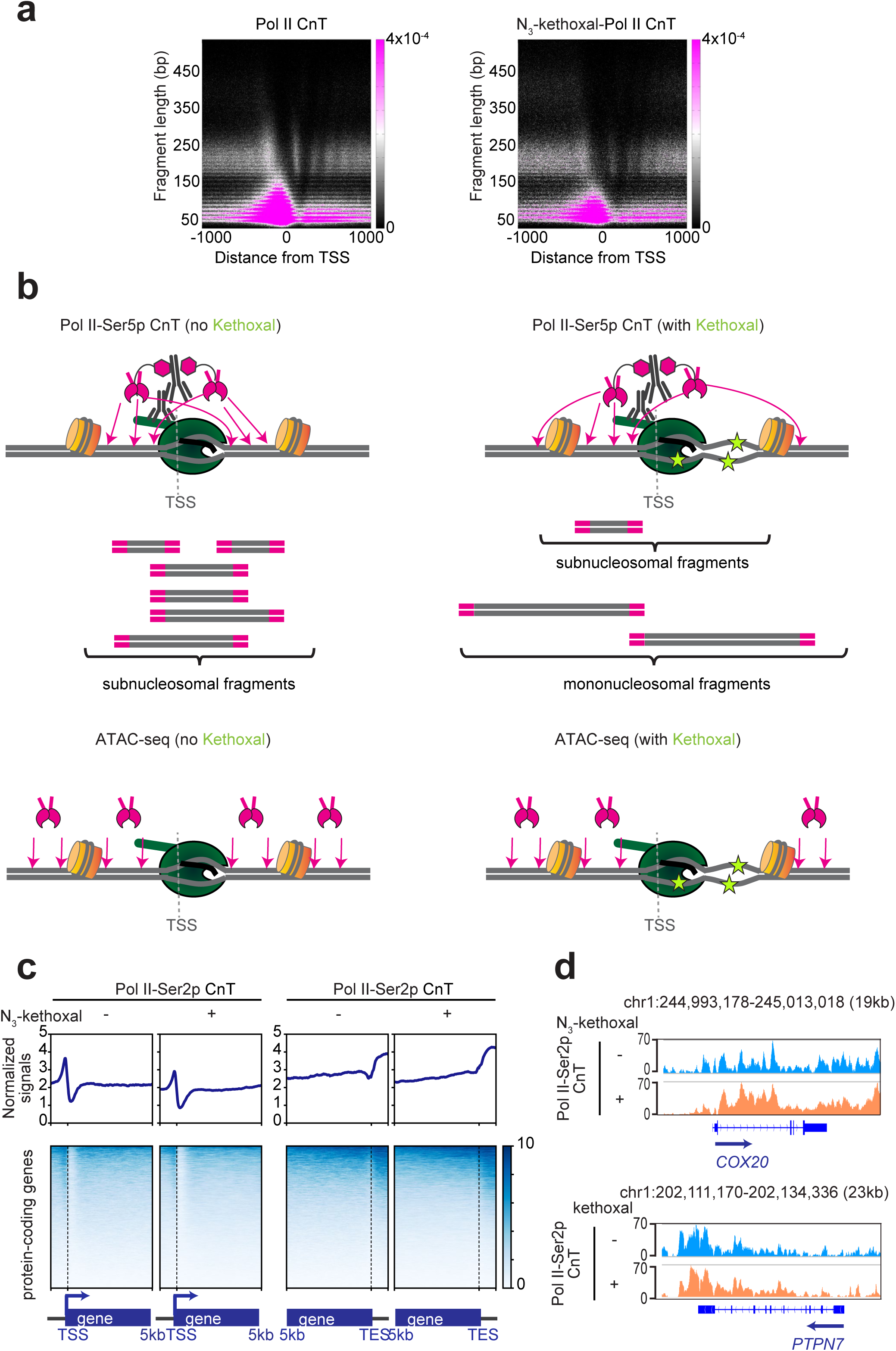
N_3_-kethoxal treatment blocks Tn5 integration into ssDNA around TSSs. a, V-plots showing fragment size versus distance from TSSs for Pol II CnT with (right) or without (left) N_3_-kethoxal treatment. The KAS-CnT total library represents N_3_-kethoxal-treated Pol II CnT. b, Schematic illustrating how N3-kethoxal blocks Tn5 integration into ssDNA between the TSS and the +1 nucleosome, causing loss of subnucleosomal fragments while preserving longer fragments derived from adjacent dsDNA in CUT&Tag and ATAC-seq. c, Average profiles (top) and heatmaps (bottom) centered on the TSSs and TESs of 12,397 protein-coding genes for Pol II-Ser2p CnT with or without N_3_-kethoxal treatment in K562 cells. KAS-CnT total corresponds to N_3_-kethoxal-treated Pol II-Ser2p CnT. d, Coverage-normalized Pol II-Ser2p CnT signals with or without N3-kethoxal in K562 cells.

**Extended Data Fig. 4.**
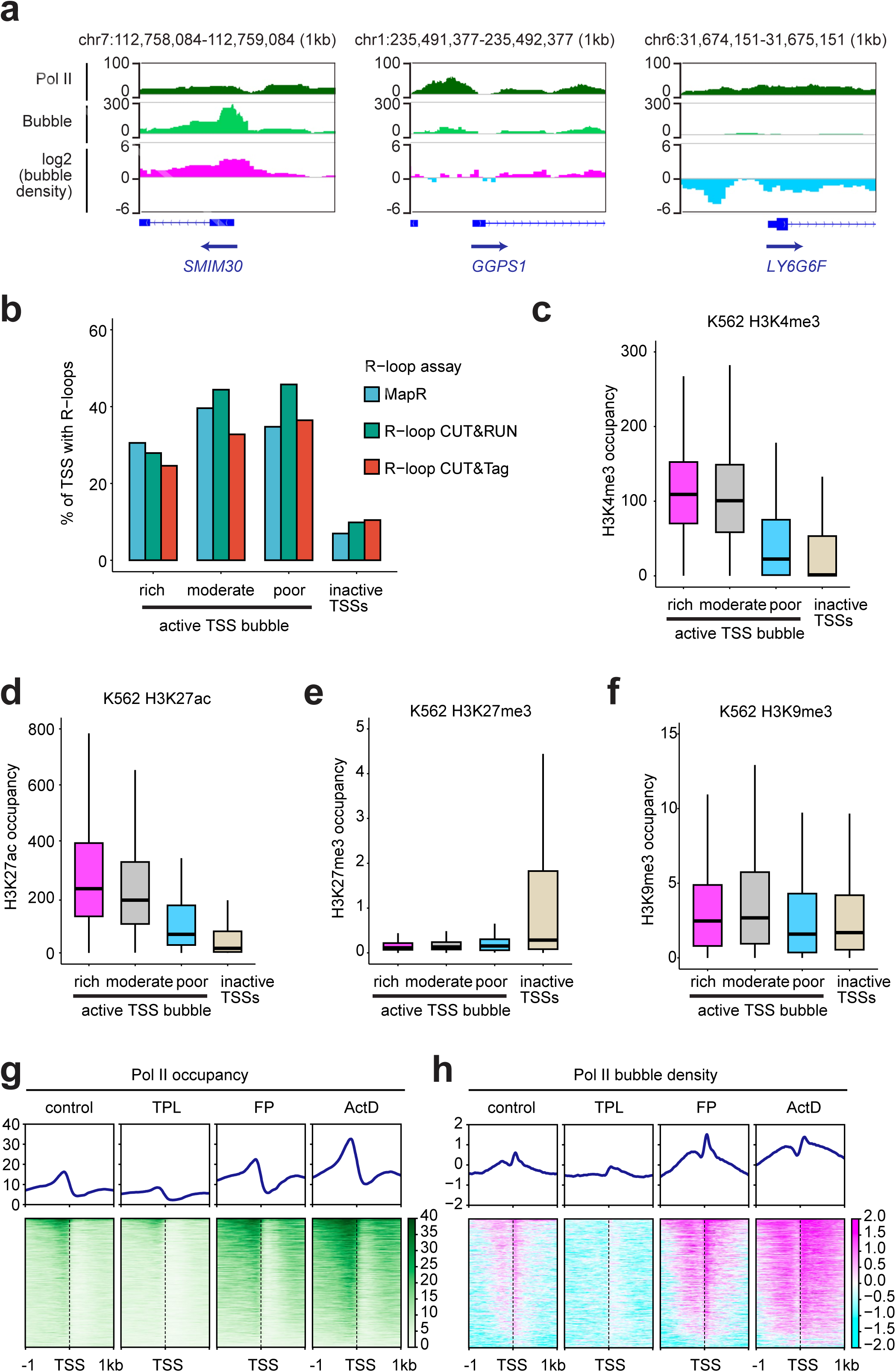
Promoter-proximal pausing increases bubble density around TSSs. a, Coverage-normalized Pol II KAS-CnT total signal, Pol II KAS-CnT IP signal corresponding to transcription bubbles and log₂(bubble density) calculated as log₂(IP/total) in K562 cells. b, Percentage of TSSs containing R-loops detected by three R-loop profiling methods in HEK293T cells. For each method, high-confidence R-loop-positive TSSs were defined as those ranking in the top 20% by R-loop signal across all replicates. c–f, Coverage-normalized H3K4me3 CnR (c), H3K27ac CnT (d), H3K27me3 CnR (e), and H3K9me3 CnT (f) signals at inactive TSSs and active TSSs classified as bubble-rich, bubble-moderate, or bubble-poor in K562 cells. g, h, Average profiles (top) and heatmaps (bottom) aligned to the TSSs of 3,859 active genes in K562 under the indicated treatments, showing Pol II occupancy (g) or log₂(Pol II bubble density) (h). Pol II occupancy is quantified by Pol II KAS-CnT total signal, and log₂(Pol II bubble density) by log₂(IP/total). Each row represents one gene.

**Extended Data Fig. 5.**
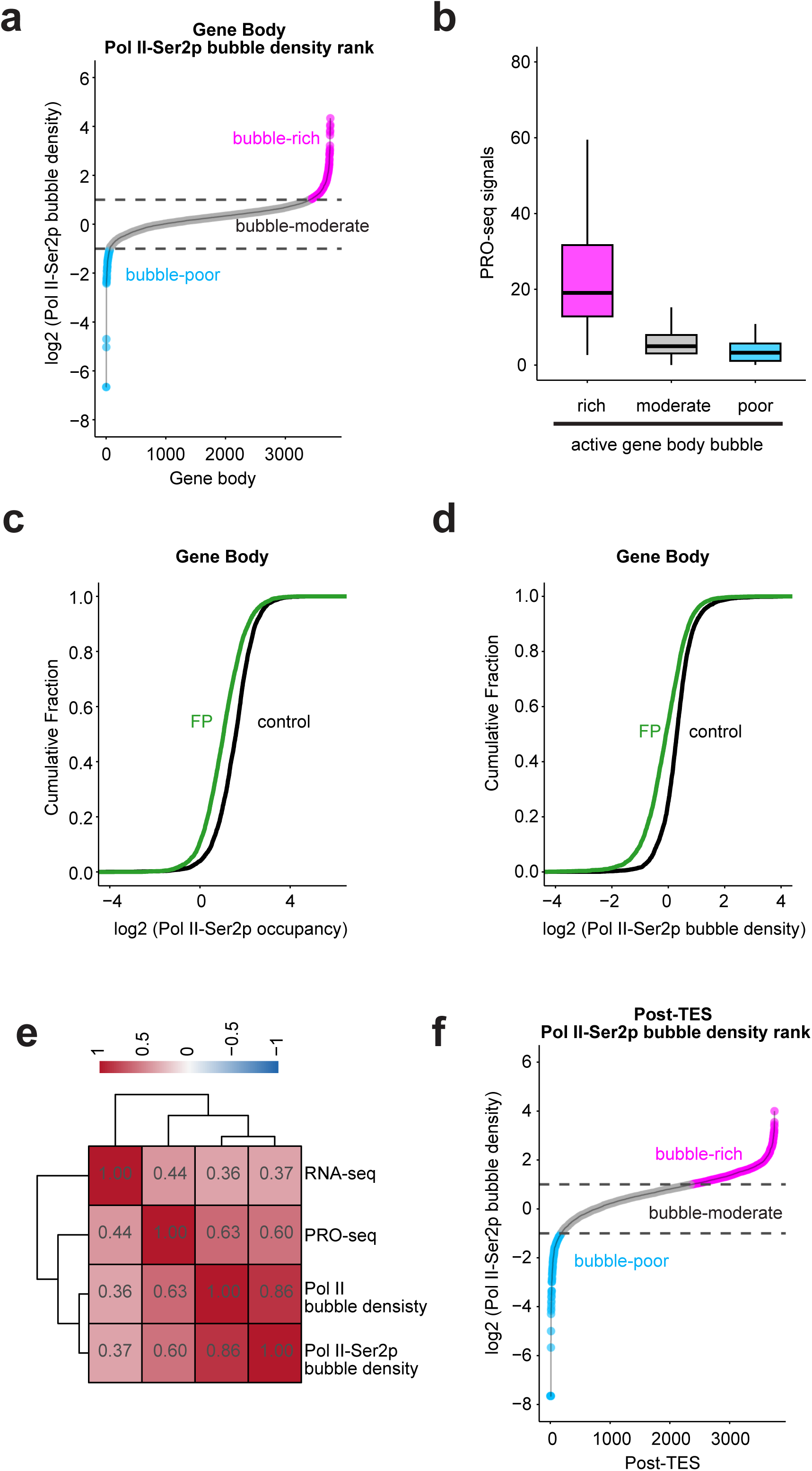
Bubble density varies across transcribed regions. a, f, Ranking of 3,754 active genes by transcription bubble density across gene bodies (a) or post-TES regions (f) in K562 cells. b, PRO-seq signals at active genes with bubble-rich, bubble-moderate, or bubble-poor gene bodies in K562 cells. c, d, Cumulative plots showing Pol II-Ser2p occupancy (c) or bubble density (d) across gene bodies in K562 cells under the indicated treatments. e, Pearson correlations among Pol II bubble density (Pol II IP/total), Pol II-Ser2P bubble density (Pol II-Ser2P IP/total), total RNA-seq signal, and PRO-seq signal across active genes in K562 cells.

**Extended Data Fig. 6.**
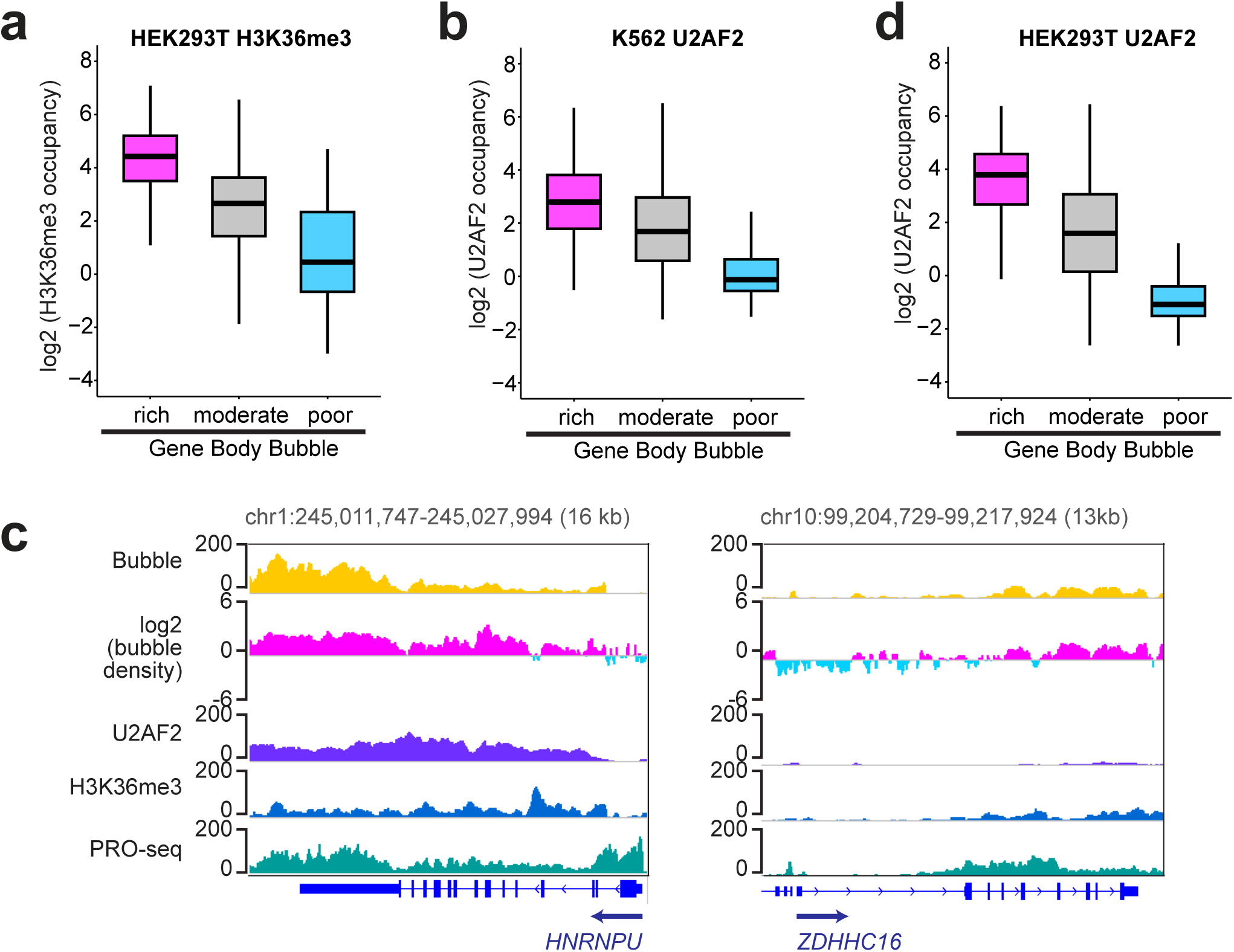
Elongating Pol II bubbles are associated with H3K36me3 and U2AF2 binding. a, Coverage-normalized H3K36me3 CnT signals across bubble-rich, bubble-moderate, and bubble-poor gene bodies in HEK293T cells. b,d, Coverage-normalized U2AF2 CnR signals across bubble-rich, bubble-moderate, and bubble-poor gene bodies in K562 (b) and HEK293T (d) cells. c, Coverage-normalized Pol II-Ser2p KAS-CnT IP, U2AF2 CnR, H3K36me3 CnT, and PRO-seq signals, together with log₂(Pol II-Ser2p bubble density), calculated as log₂(IP/total), in HEK293T cells.

**Extended Data Fig. 7.**
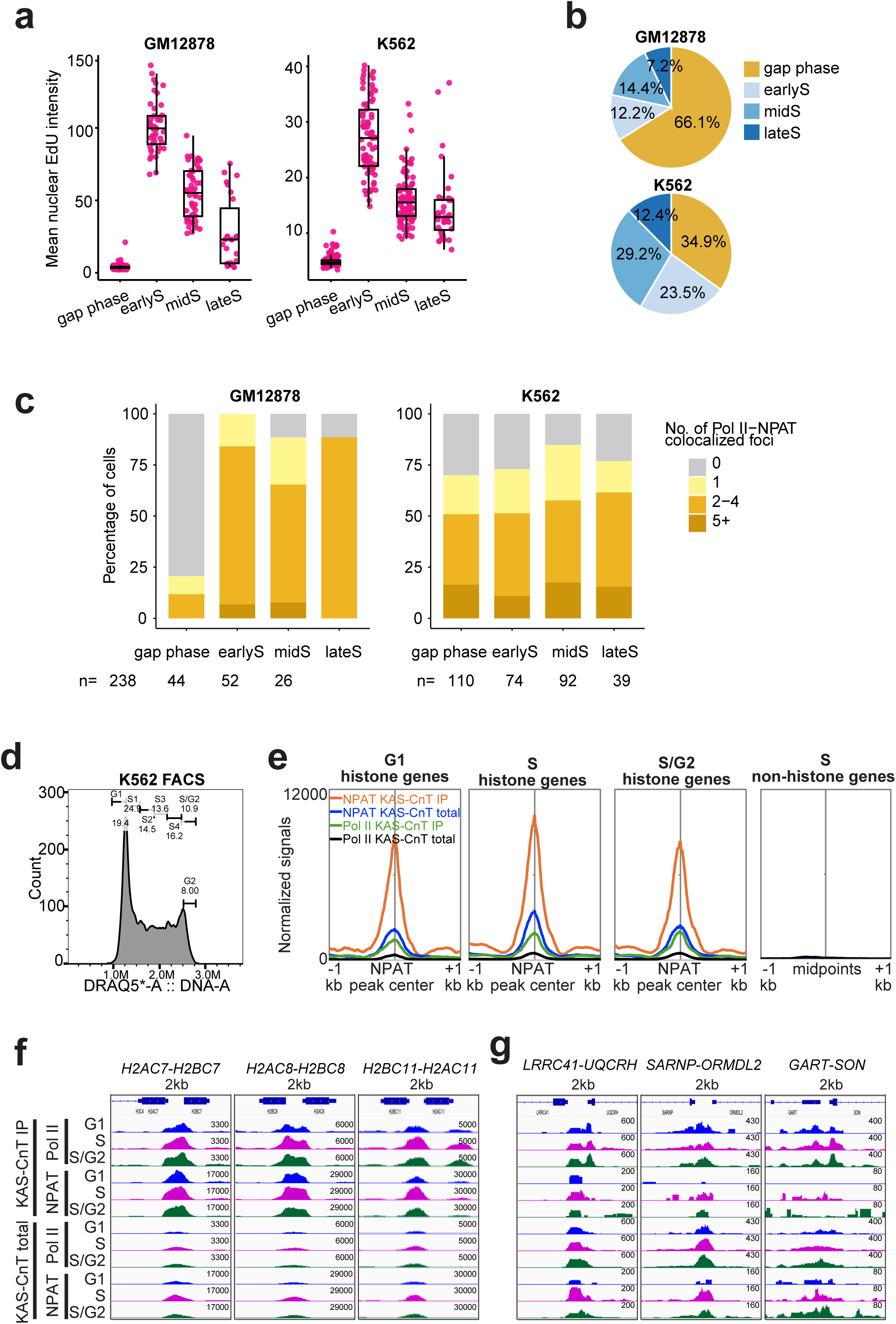
Dense transcription bubbles persist at RC histone genes throughout the cell cycle in K562 cells. a, Mean EdU intensity in GM12878 and K562 nuclei at the indicated cell-cycle stages. EdU staining patterns distinguished early, mid, and late S phase, while EdU-negative interphase cells were classified as gap phase cells. b, Percentage of cells in each cell-cycle stage in GM12878 and K562. c, Distribution of GM12878 and K562 cells across the indicated cell-cycle stages by number of NPAT–Pol II-Ser5p colocalized foci. N indicates the number of cells analyzed at each stage. d, Isolation of cell cycle fractions. FACS gating schema to purify G1, S-phase, and G2 populations of K562 cells. Even-width gates were fit to the distribution of DNA content in exponentially proliferating K562 cells. Mitotic cells were gated out based on DRAQ5 eccentricity. Percent of the parent population is shown for each gate. G1 cells were defined as left of the first mode. S-phase cells were collected from the gate labeled S3. S/G2 cells were collected from the last gate. Post-hoc analysis by fitting a gate right of the second mode (labeled G2 only) suggested that 8.0% of cells are in G2-phase, indicating 73% G2 purity in the S/G2 population. e, Average plots centered on 36 NPAT KAS-CnT peaks at the 5′ ends of RC histone genes, showing Pol II and NPAT KAS-CnT IP and total signals in G1, S, and S/G2 phase FACS-isolated K562 cells. An average plot centered on the midpoints of 12 bidirectional non-histone gene pairs in S phase cells is shown on the same scale to indicate background signals. f,g Coverage-normalized signals for Pol II and NPAT KAS-CnT IP and total at representative bidirectional H2A/H2B gene pairs (f) and bidirectional non-histone gene pairs (g) in K562 cells.

**Extended Data Fig. 8.**
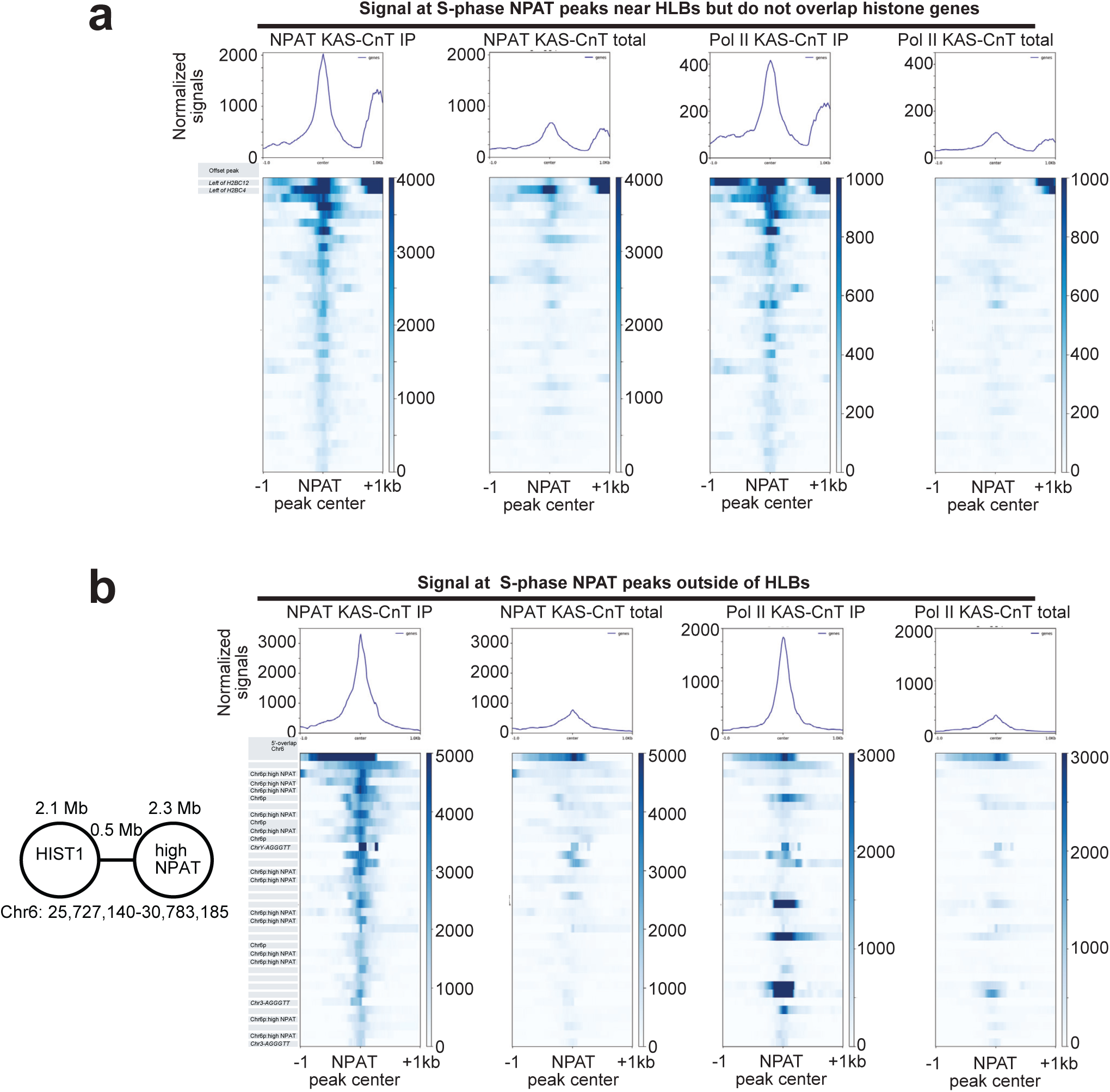
NPAT is juxtaposed to transcription bubbles within or near HLBs. a, Average profiles (top) and heatmaps (bottom) centered on the top 36 NPAT peaks (based on NPAT KAS-CnT IP signal) located within HLBs that do not overlap RC histone genes in K562 S-phase cells. b, Same as (a), but for the top 36 NPAT peaks located outside of HLBs.

